# Neural Correlates of Temporal Anticipation

**DOI:** 10.1101/2022.12.15.520557

**Authors:** Matthias Grabenhorst, Georgios Michalareas

**Affiliations:** Ernst-Strüngmann Institute for Neuroscience in Cooperation with Max Planck Society, Deutschordenstr. 46 60528 Frankfurt am Main; Max Planck Institute for Empirical Aesthetics, Grüneburgweg 14, 60322 Frankfurt am Main

## Abstract

Temporal anticipation is a fundamental process underlying complex neural functions such as associative learning, decision-making, and motor-preparation. Here we study event anticipation in its simplest form in human participants using magnetoencephalography. We distributed events in time according to different probability density functions and presented the stimuli separately in two different sensory modalities. We found that the temporal dynamics in right parietal cortex correlate with reaction times to anticipated events. Specifically, *after* an event occurred, event probability was represented in right parietal activity, hinting at a functional role of event-related potential component P300 in temporal expectancy. The results are consistent across both visual and auditory modalities. The right parietal cortex seems to play a central role in the processing of event probability density. Overall, this work contributes to the understanding of the neural processes involved in the anticipation of events in time.

## Introduction

The anticipation of future events is an elementary function of neural systems that is crucial to the survival of an organism. At the scale of seconds, an animal needs to predict a predator’s immediate actions in order to evade its attack. In humans, temporal prediction at short time scales underlies many different aspects of cognition in a wide variety of tasks, including in sports, music making, and everyday choice behavior. Consequently, the identification of neural correlates of predictive processes has been a focus of recent work. A considerable part of this work in non-human primates^1-4^ and humans^5-10^ builds on the hazard rate (HR) as a canonical model of anticipatory neural activity^11^. Initially, the prominent HR was proposed as a model of reaction time (RT) behavior^12^ before it was used as a model of neural anticipation correlates. HR represents the probability that a future event is imminent, given that it has not already occurred^12,13^. Based on this intuitive computational assumption, neural activity in anticipatory tasks was described at different levels ranging from single cells to population responses.

In non-human primates, single neurons’ firing rates were correlated with the HR in visual areas V1^4^ and V4^1^, and in lateral parietal area LIP^2^, suggesting that this model of anticipation is implemented in brain activity from early sensory to later stages of information processing. In humans, the HR was proposed as a model of population responses in temporal anticipation, based on correlations between HR and theta (4–8 Hz)^6^, beta (13–30 Hz)^9,10^, and gamma (40–70 Hz)^9^ band neural oscillations, as well as modulation of gamma band coherence between motor cortex and spinal cord neurons^9^. In work using fMRI, where experimental designs are tailored to the BOLD response resulting in tasks that cover longer time spans in the seconds range^7,14^, correlates of the HR computation were proposed in primary visual cortex and extra-striate areas V2/V3^5^ as well as post-stimulus BOLD modulation according to the cumulative HR in supplementary motor area and superior temporal gyrus^7^. Taken together, previous work proposes a variety of neural HR correlates in diverse areas ranging from early to later stages in the cortical hierarchy and in both activity preceding and following an event in time.

Most of the work on the HR employs a psychometric-neurometric mapping approach in a wider sense. This strategy builds on models of anticipatory behavior to inform analysis of neural data. However, recent work found the HR to be an inadequate model of behavior and thereby challenged the HR as a canonical neural computation in event anticipation^15,16^. This work featured simple experimental tasks that are highly similar to those used in work promoting the HR as a model of both, RTs and the neural processes that generate them. This work further questions the HR’s biological plausibility because of its computational complexity^13^ and numeric instability^17^, and also because of the HR’s immense range: in the commonly used cases of uniform and Gaussian event distributions with event certainty (P_event_ = 1), the HR increases with a very steep slope towards the upper limit of the time span over which it is computed. It is unclear whether the brain can compute such steep slopes over short time spans. This mathematical property is sometimes addressed by truncation of the HR variable to avoid the impractical model prediction of positive infinity. In contrast, an alternative model of anticipatory behavior implies that the brain computes the reciprocal event probability density function (1/PDF) as its model of events distributed in time^15,16^. The 1/PDF model is computationally parsimoneous as the estimation of PDF can be approximated by the relatively simple operations of event counting and time estimation.

Apart from HR-based work, the effect of probability on anticipatory neural processes is commonly investigated in activity that is time-locked to external events and also in the time-frequency representation of neural data. There is a vast literature on the effects of probability on event related potentials (ERP). Classic examples include the P300^18^, a component whose ERP amplitude is inversely related to target probability^19-21^. Interestingly, the P300 may also be modulated by temporal probability, as suggested by work manipulating the number of items of a category over time^22^. In tasks with a simple temporal structure, the P300 co-varies with expectancy, driven by e.g. two different stimulus onset times^23^. Still the effects of event probability density on the component are unclear. The posterior N2 (N2c) is another ERP component that is sensitive to target probability^24^ and is associated with stimulus categorization. These two late components often occur *after* a response was given^25^ which relates these findings less readily to anticipatory processes. Examples of *pre-stimulus* ERP components include the stimulus-preceding negativity (SPN)^26^, a signal argued to reflect stimulus anticipation, the lateralized readiness potential (LRP)^27^, a preparatory motor signal predictive of response times, and the contingent negative variation (CNV)^28,29^. The CNV is linked to timing processes^30^ and is proposed to reflect temporal expectancy and response preparation and originates in motor-related areas^31-34^. The CNV is suggested to co-vary with the HR^6^.

Guided by the neural correlates of event anticipation and based on our recent behavioral work that challenges the HR model, we investigate the neural processing of events distributed stochastically in time. This paper addresses several aims.

First, we test whether human anticipatory RT behavior is adequately described by the HR model or by the 1/PDF model. Both models are based on event probability over time. They are not mutually exclusive but differ in their computational assumptions^16^.

Second, based on the results of the first analysis, we investigate how event probability over time affects neural activity that is time-locked to sensory events. We investigate potential modulation of pre-stimulus time-locked activity, as well as early and late components of event-related fields (ERF) as measured with magnetoencephalography (MEG). We aim to localize the cortical origin of ERF components of interest.

All experiments and analyses are performed separately in two sensory modalities to identify anticipatory neural activity that is shared by vision and audition and may therefore be independent of modality-specific processes.

## Results

### Set–go experiment and probabilistic design

The participants performed a ‘set’–’go’ task while neural activity was recorded with MEG. The task, which consisted of separate auditory and visual experimental blocks, demanded the participants to respond as quickly as possible to the ‘go’ cue with a button press (**Fig. 1a**). In trials without a ‘go’ cue (catch trials), participants were asked to not press the button. The time span between ‘set’ and ‘go’ cues was a random variable determined by either an exponential (‘Exp’) or a flipped exponential distribution (‘Flip’). Each of these distributions was defined over timespan t = [0.4, 1.4] s and was fixed per experimental block (**Methods)**. A rapid response to the ‘go’ cue required participants to form predictions about the temporal-probabilistic structure embedded in the task, with the assumption that better temporal prediction is reflected in faster reaction times.

**Fig. 1.**
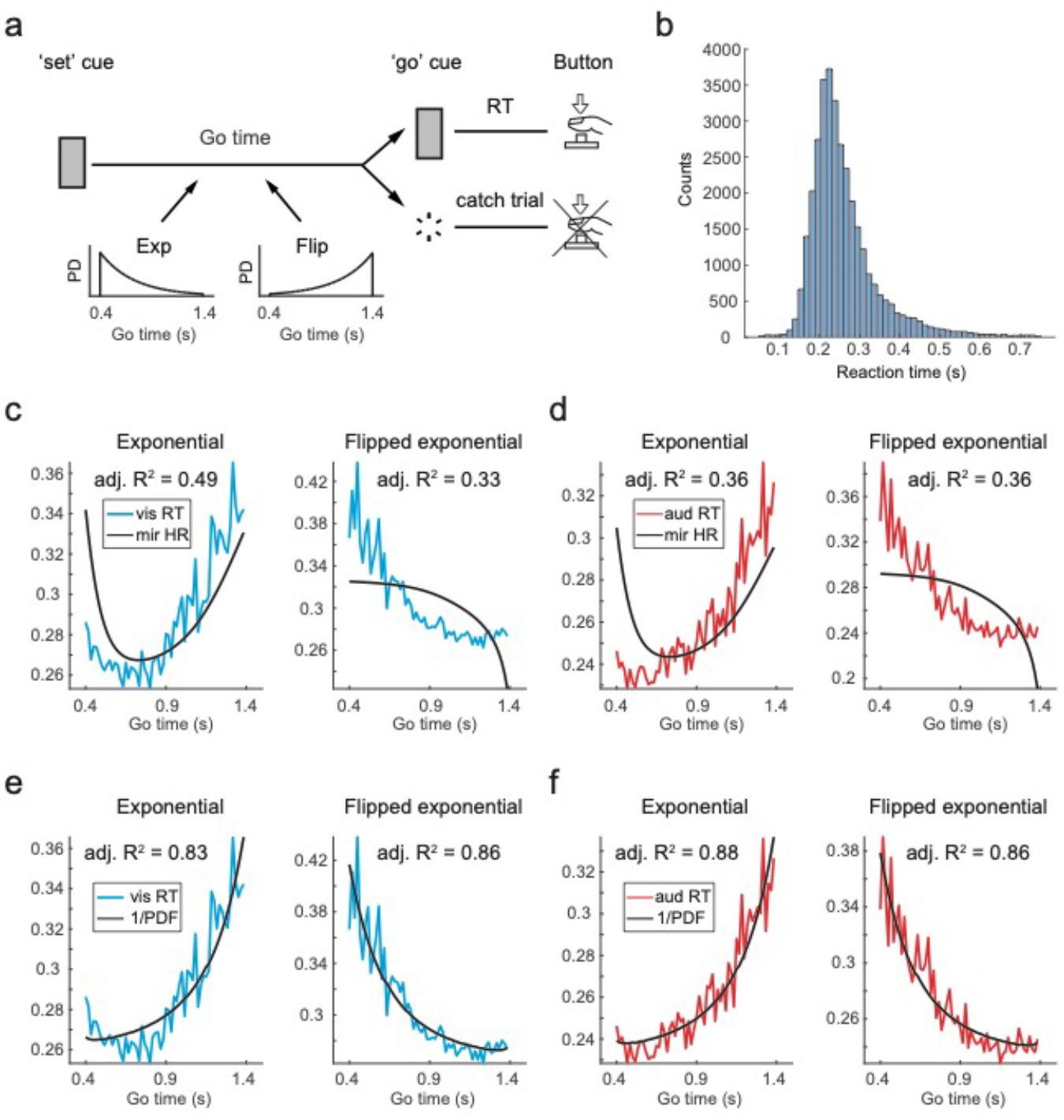
Task, reaction time (RT) data and model fits. **a**) Schema of visual and auditory task. Time between ‘Set’ and ‘Go’ cues (white noise bursts in audition and checker-boards in vision, both 50 ms in duration) was drawn from exponential and flipped exponential probability density (‘PD’) functions, both defined over t = [0.4, 1.4] s, in a block-wise design (**Methods**). Participants were asked to respond as fast as possible to the ‘go’ cue by pressing the button. In absence of ‘go’ cue (catch trial) no button should be pressed. **b**) Histogram of RT pooled across conditions and subjects from all analysed trials (N = 33354). **c**) Mirrored temporally blurred hazard rate (HR) fit to RT from the visual exponential (left) and flipped exponential (right) conditions. **d**) Mirrored temporally blurred HR fit to RT from the auditory exponential (left) and flipped exponential (right) conditions. **e**) Reciprocal probabilistically blurred PDF fit to visual RT and **f**) auditory RT. All shown RTs correspond to the trials that were selected for MEG analysis.

In analysis, we tested the canonical model of event anticipation, the HR-based model of RT, and the recently proposed PDF-based model as candidates of the participants’ representation of event probability density.

### Reciprocal event PDF captures reaction time modulation

24 participants generated a total of 33354 RTs (**Fig. 1b**). In all four conditions, the shape of RT distributions was similar with a steep left flank and a right-skew (**Fig. S1**), resembling classic RT distributions from simple RT tasks^12^. Average RT was shorter in audition than in vision in both exponential (∆ _RT_ = - 28.4 ± 5.8 ms, mean ± SEM, *P* = 0.02, *t*_*23*_ = - 2.42 ) and flipped exponential conditions (∆ _RT_ = - 32.5 ± 8.0 ms, mean ± SEM, *P* = 0.009, *t*_*23*_ = - 2.75 ). In order to investigate the influence of event probability density on RT, mean RT was plotted over ‘go’ time (**Fig. 1c**). The shape of the RT curves suggested an inverse relation to the event PDF: where event PDF was large, RT was small and vice versa. Note that near the extremum of the ‘go’ time span where event probability is highest, i.e. at ‘go’ time = 0.4 s for exponential and 1.4 s for flipped exponential distributions, the RT curves do not monotonically decrease but increase instead. This phenomenon is most evident in the exponential case and is more pronounced in vision than in audition. Apart from this modality-driven effect, the RT modulation was highly similar in vision and audition which indicates that participants extracted similar temporal-probabilistic information embedded in the task, independent of input modality.

In order to investigate these qualitative findings further, we tested how well the RT curves can be fit by two models, one based on HR and one based on PDF itself. Specifically, in the first model, we fit as explanatory variable the mirrored temporally blurred HR (**Methods**) to the RT curves (**Fig. 1c** and **d**). In the exponential case, the HR model did not provide an acceptable model fit as it strongly over-estimated RT at short ‘go’ times. In the flipped exponential condition, the HR model was even qualitatively inadequate: the RT curve decreased in a concave way, yet the HR model predicted a convex RT curve. These findings are reflected by small values of adjusted R^2^. In the second model of RT, we used as explanatory variable the reciprocal probabilistically blurred PDF to RT. In both exponential and flipped exponential conditions and in both modalities, the model adequately captured RT modulation (**Fig. 1e** and **f**). The large values of adjusted R^2^ and the absence of systematic deviations between data and fit line indicate a remarkably good fit given such a simple descriptive model.

In sum, the reciprocal probabilistically blurred event PDF captured well the RT modulation, while the canonical HR-based model failed to do so. These behavioral results replicate our previous work^15,16^ and provide us with a model of RT that is used in the analysis of neural data.

### MEG data preprocessing

All MEG data analysis was performed in MatLab (The MathWorks Inc., Natick, USA) using the FieldTrip toolbox for EEG/MEG analysis developed at the Donders Institute for Brain, Cognition, and Behavior (Nijmegen, The Netherlands)^35^. Artifactual data segments that contained jumps in the SQUIDs (super-conducting quantum interference devices), eye movements, heart and muscle activity were identified and removed using semi-automated FieldTrip routines (**Methods**).

### Event-related field components

We investigated potential effects of event probability density on neural activity time-locked to stimulus events. First some basic sanity checks were performed. At the single-trial level, within-participant, the MEG data was split into three sets and time-locked relative to the onset of ‘set’ and ‘go’ cues and to the time point of the button press. The simple ‘set’ and ‘go’ stimuli resulted in a prominent M100 (P1) wave in both vision (**Fig. 2a** and **b**, top row) and audition (**Fig. 2d** and **e**, top row). The distribution of activity over the sensor array during the time span from 55 to 85 ms post cue indicates visual (**Fig. 2a** and **b**, bottom row) and auditory (**Fig. 2d** and **e**, bottom row) sensory cortex as the likely source of activity for data locked to ‘set’ and ‘go’ cues. The location of the ERF related to the button press (**Fig. 2c** and **f**, top row) roughly corresponds to left motor cortex (**Fig. 2c** and **f**, bottom row) which is in line with participants pressing the button with their right index finger. Note that location of activity on the sensor array does not correspond to location on the human brain surface since participants differed in head size and in the positions they take relative to the MEG sensor array. At this level of grand averages, the time course of early neural activity is quite similar *across* the Exp and Flip probability conditions in all three different data sets (locked on ‘set’ and ‘go’ cues and on button) and in both vision (**Figs. 2a** and **b**, both left vs. right) and audition (**Figs. 2d** and **e**, both left vs. right). Nonetheless, there are differences in the time course of the ERFs *across* the ‘set’ and ‘go’ cues *within* each of the Exp and Flip conditions. These differences are more pronounced in later ERF components which is to be expected since the button press occurs around 200 ms after the ‘go’ cue (but not after the ‘set’ cue). We investigate the relationship between these components and event probability density below.

**Fig. 2.**
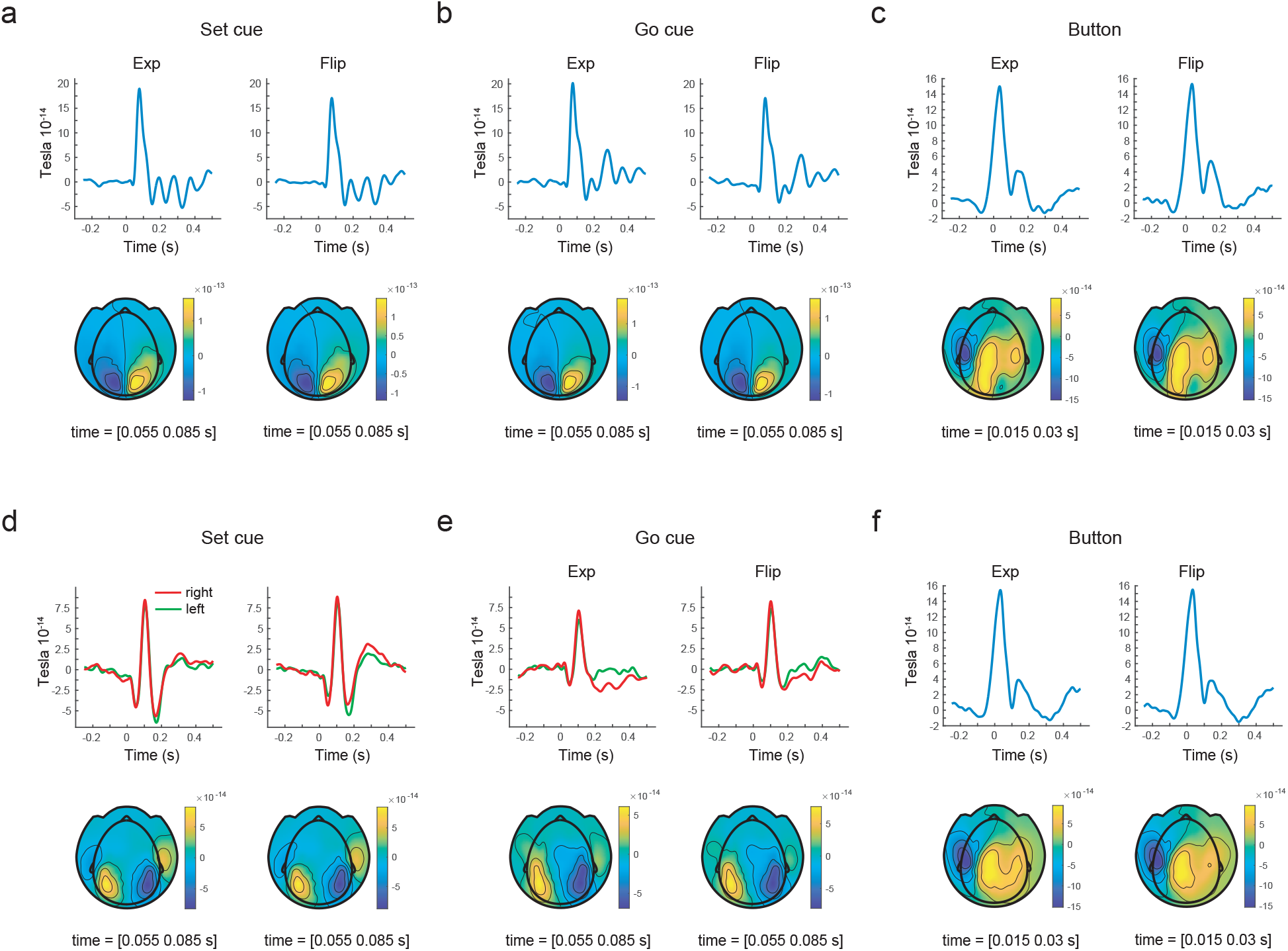
Event related fields evoked by visual and auditory stimulation and by the button press. ERFs were averaged across all channels that comprise the positive part of a dipole selected at the single-subject, within-condition level (**Methods**).

### Event-related field components – correlation-based analysis strategy

The first behavioral analysis has shown that the event PDF is related to RT in a systematic way, with short RTs occurring when PDF is large and vice versa. This suggests that the brain anticipates the ‘go’ cue by computing an estimate of the event PDF. We therefore focused on the epoch of the ‘go’ cue in order to investigate potential neural signatures of both the anticipatory process, *before* the ‘go’ cue, and the post-stimulus processes, *after* the ‘go’ cue. We employed a correlation-based approach that related RT to neural activity time-locked to the ‘go’ cue. Specifically, we were interested in pre-’go’ cue activity as a potential correlate of preparatory activity as well as in early and late ERF components. The analysis comprised of three steps (**Methods**):

1) In each of the four experimental conditions, at the single-participant level, the MEG data was aggregated within adjacent pairs of consecutive ‘go’ times (these pairs are called *frames* from this point on). The aggregation reduced the number of unique ‘go’ times from 60 to 30 (= 30 frames). This mild smoothing was performed to reduce noise in the data before correlation. Although the frames differ in the number of trials, this averaging did not introduce a bias in the mean within each bin due to the close to Gaussian distribution of the ERF data and thus no bias in the correlation (**Supplementary Methods**).

2) At the single-participant level, for each channel-by-time-point duplet, Spearman’s rho was computed between the 30 frames of MEG data and the 30 averaged RTs. Note that the resultant rho has the same dimensionality as the within-participant grand average of MEG data (channels-by-time-points).

3) On all participants’ rho, a cluster-based permutation test was run to identify channel-by-time clusters in which rho differs from zero, indicating an interpretable relationship between MEG data and RT.

### Event-related field components – correlation with RT at the sensor level

First, we aimed to identify significant correlation clusters between the ERF and RT at the sensor-level. Across all four conditions, the cluster-based permutation test revealed a clear pattern of clusters, i.e. of channel-by-time duplets, where rho differed significantly from zero (see **Tbl. S1** for all clusters’ *P* values). A positive cluster was identified around 200 ms (**Fig. 3a**) after ‘go’ onset indicating that the differences in (positive) rho from zero was most pronounced over left lateral sensors (**Fig. 3c**, left). As could be expected, given that the correlation was performed on dipolar scalp topography, a corresponding negative cluster was identified also around 200 ms post ‘go’ (**Fig. 3b**) which included mostly right centro-occipital sensors (**Fig. 3d**, left). The interpretation of this positive/negative cluster pair as reflecting motor cortex activity is supported by the distribution of RT (**Fig. S1**): since RTs are distributed in time relative to the ‘go’ cue, the correlation-based analysis should identify a significant correlation between RT and neural activity in motor cortex. Although the location of this dipole (**Fig. S2a**, left, and **b**, left) may suggest left motor cortex as the source of neural activity, care must be taken since the sensor-level topography might be a superposition of multiple sources with some located in the right hemisphere.

**Fig. 3.**
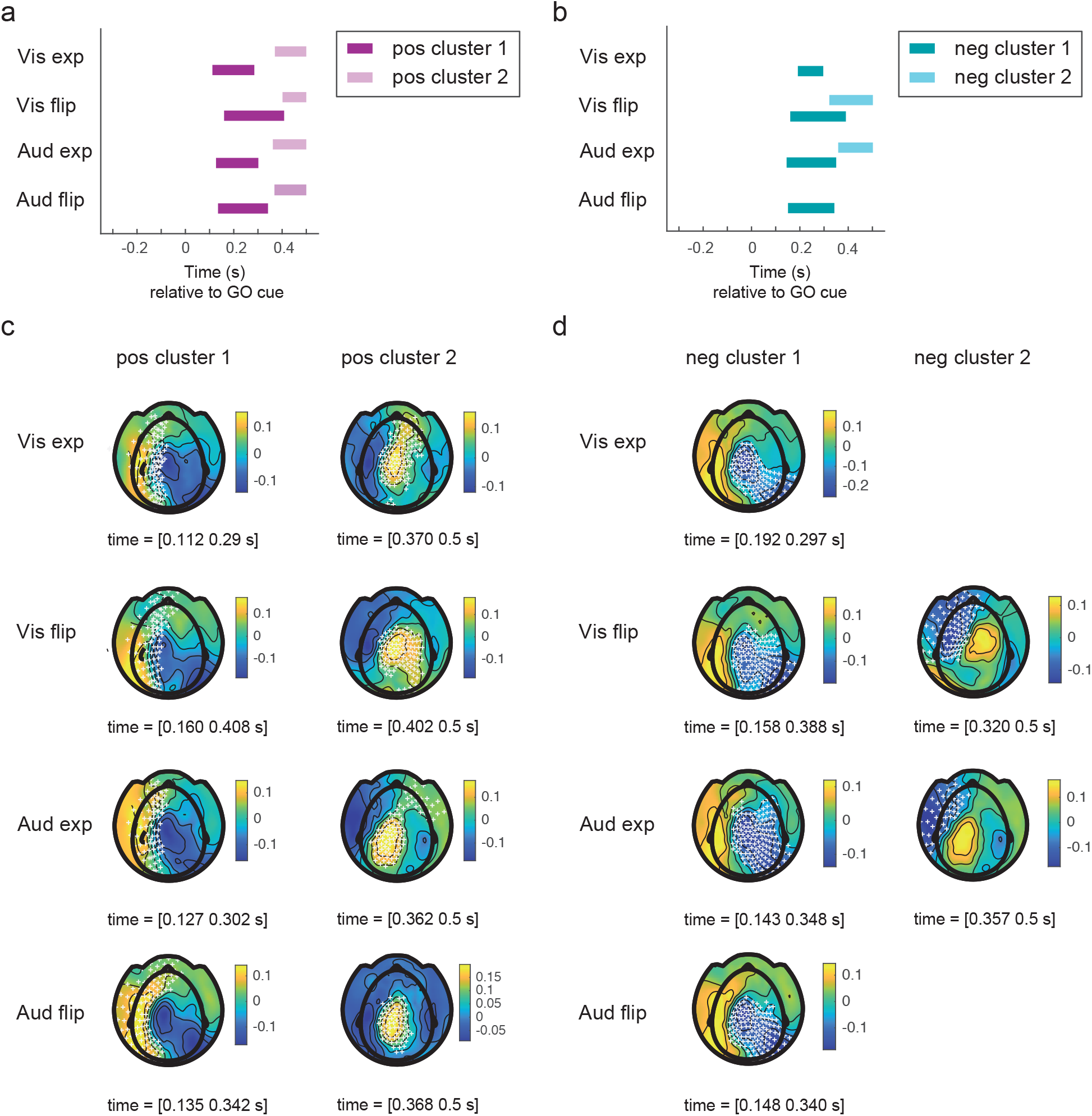
Cluster-based permutation test on Spearman’s rho (correlation between ‘go’ cue ERF and RT). **a**) Positive clusters. **b**) Negative clusters. **c**) Topography plots of rho, channels of positive cluster highlighted. **d**) Topography plots of rho, channels of negative cluster highlighted. A minimum time span of 20 ms was set for a channel to be included in a cluster.

Second, around 400 ms post ‘go’ cue, a positive cluster was identified (**Fig. 3a**), corresponding to central sensors (**Fig. 3c**, right). In two of the four conditions, a corresponding negative cluster was found (**Fig. 3b**), whose topography comprised of left latero-frontal sensors (**Fig. 3d**, right). Note that these late clusters’ topography roughly resembles the inverse polarity of the early cluster around 200 ms (compare **Fig. 3c**, left with **d**, left, and **c**, right with **d**, right). Again, a definitive source of this signal is not readily apparent due to the broad distribution over the sensor array that obscures whether the dipole (**Fig. S2a**, right, and **b**, right) is indeed restricted to one hemisphere.

As a last step in the sensor-level analysis of evoked activity, we investigated the pre-’go’ cue time span. This correlation-based analysis is identical to the one above, with the exception that the MEG data was baseline-corrected relative to a pre-’set’ cue period (**Methods**) in order to investigate potential anticipatory activity reflected in the ERF prior to the ‘go’ cue. For RT as regressor, the same clusters were identified as in the analysis above, where a pre-’go’ baseline was used. Importantly, no cluster was identified during the pre-’go’ cue time span indicating that time-locked activity preceding the ‘go’ cue did not significantly correlate with RT (**Tbl. S2** and **Figs. S3** and **S4**). Thus, our analysis did not reveal any neural activity before the onset of the ‘go’ cue that is correlated with RT. This indicates that at the level of activity time-locked to the ‘go’ cue, no preparatory activity could be identified.

Taken together, this first correlation analysis between sensor-level evoked brain activity and RT showed that there are two epochs where they are significantly related, an early one around 200 ms and a late one around 400 ms after the ‘go cue’. In the first case, the analysis identified clusters around 200 ms post ‘go’ cue, which was expected since participants pressed the button around this time point (**Fig. 1b**). This was also reflected in the distribution of these early clusters on the sensor array, suggesting a source in left motor cortex (button pressed by right hand index finger). In the second case, the analysis identified late clusters around 400 ms post-’go’ cue, a time span associated with the probability-sensitive P300 wave^19-21,31^.

### Event-related field components – correlation with RT at the source level

We aimed to identify source-level representations of the correlation results described above at the sensor-level to obtain a more precise spatial estimate of the neural activity correlated with RT which may be informative about the potential role of these ERF components in the processing of event probability density. As was the case at the sensor-level, the single-trial ERF data relative to the ‘go’ cue was aggregated in frames (see above). The per-frame data were then projected into source-space and Spearman’s rho was computed between source-space ERF and RT for each source and time point (**Methods**). The distribution of rho was centered around zero in all four conditions, resembling a Gaussian distribution (**Fig. S5**). Rho was averaged within time windows of interest (**Tbl. S3**) that were selected based on the clusters identified in sensor-space analysis. A cluster-based permutation test was run on these aggregated rho data to identify clusters of sources in which rho differed significantly from zero.

An important limitation of the cluster-based permutation test, as currently implemented in FieldTrip, is that it cannot perform 4-D clustering, i.e. along the 3 spatial dimensions of the volumetric source space and time at once. It can only perform 3D spatial clustering in source space using a low-level function of SPM (*spm_bwlabel*.*m*) inside the function that estimates clusters (*findcluster*.*m*). Given this limitation, rather than performing such a 3D cluster permutation test for each time point and then having to correct for a large number of multiple comparisons (number of time points), we used the cluster time spans already identified in sensor-level analysis and performed a single test for the time span of each cluster. This test identified for the time period of a cluster found in sensor space, the corresponding sources that have a significant correlation with RT.

The above analysis pipeline identified patterns of positive and negative clusters around both 200 and 400 ms which were consistent across all four conditions and were significant in most (but not all) cases (**Tbl. S4**).

A striking pattern of clusters emerged, in which the early positive clusters were associated with activity in right parietal cortex and the early negative clusters with activity in left parietal and motor cortex (**Tbl. 1**). This pattern was reversed for the late clusters: positive clusters were located in left parietal and motor and negative ones in right parietal cortex. Notably, in all four conditions a second positive cluster was identified around 200 ms post-’go’ cue which was comprised of sources localized in the cerebellum.

**Table 1.**
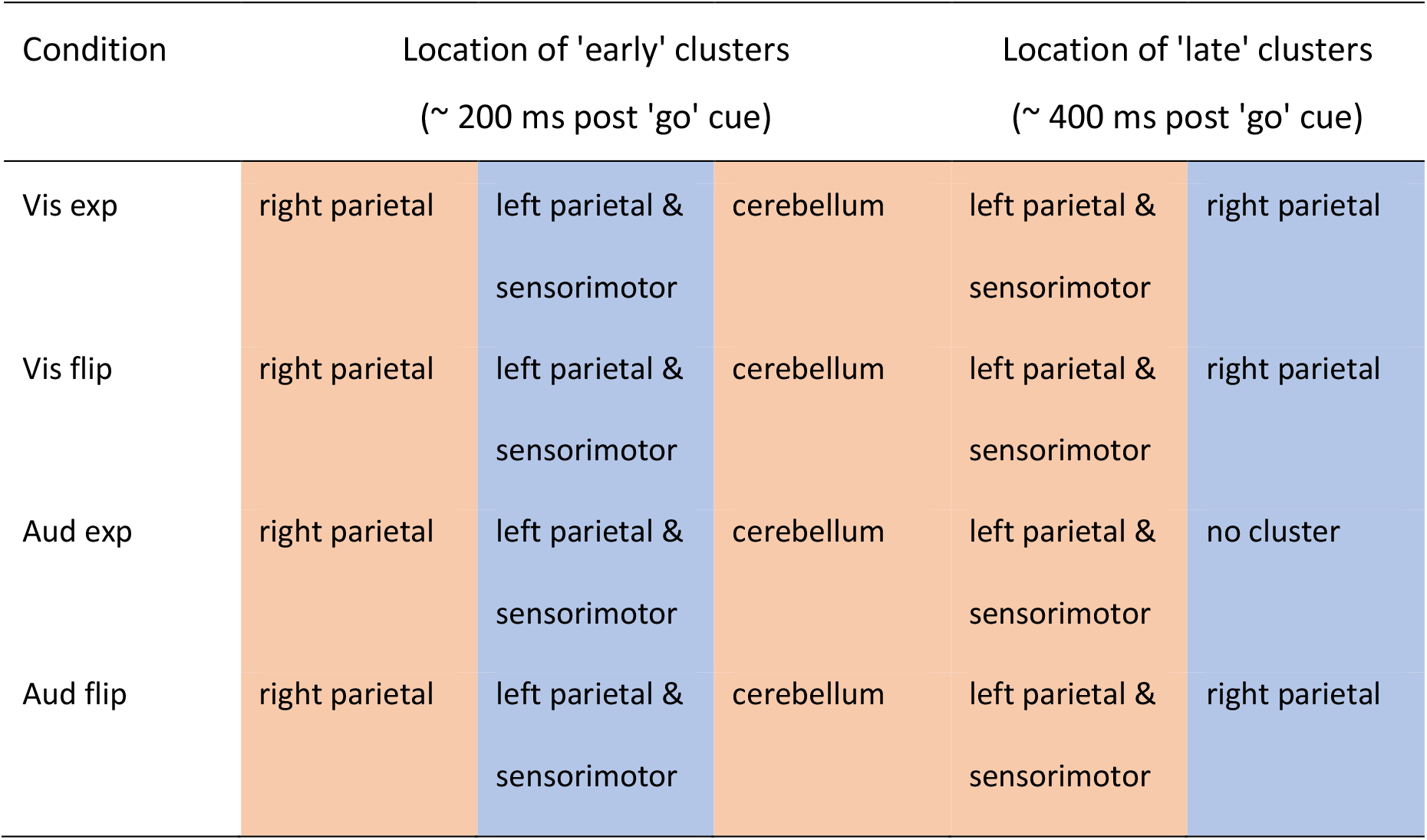
Location of source clusters computed on Spearman’s rho (correlation between ‘go’ cue ERF and RT). Positive clusters (red), negative clusters (blue).

| Condition | Location of 'early' clusters<br>(~ 200 ms post 'go' cue) |  |  | Location of 'late' clusters<br>(~ 400 ms post 'go' cue) |  |
| --- | --- | --- | --- | --- | --- |
| Vis exp | right parietal | left parietal &<br>sensorimotor | cerebellum | left parietal &<br>sensorimotor | right parietal |
| Vis flip | right parietal | left parietal &<br>sensorimotor | cerebellum | left parietal &<br>sensorimotor | right parietal |
| Aud exp | right parietal | left parietal &<br>sensorimotor | cerebellum | left parietal &<br>sensorimotor | no cluster |
| Aud flip | right parietal | left parietal &<br>sensorimotor | cerebellum | left parietal &<br>sensorimotor | right parietal |

**Fig. 4** shows the across-condition averages of the sources that comprise early and late clusters. The early *negative* clusters align well with left sensorimotor cortex, as well as with primary somatosensory cortex (**Fig. 4a**, right), likely reflecting activity related to the button press in generation of RT. This interpretation of the negative correlation between RT and ERF is supported by the fact that in simple RT tasks, RT variance is positively correlated with average RT^12^ which is also seen in the distribution of our RT data (**Fig. S1**). The right-skewed distribution of RT should lead to a larger within-frames-averaged ERF amplitude where RT variance is small (short RT) and to a smaller average ERF amplitude where RT variance is large (large RT)^25^. Therefore, the distribution of the early negative clusters over motor-related areas can be seen as a sanity check that supports our correlation-based analysis strategy.

**Fig. 4.**
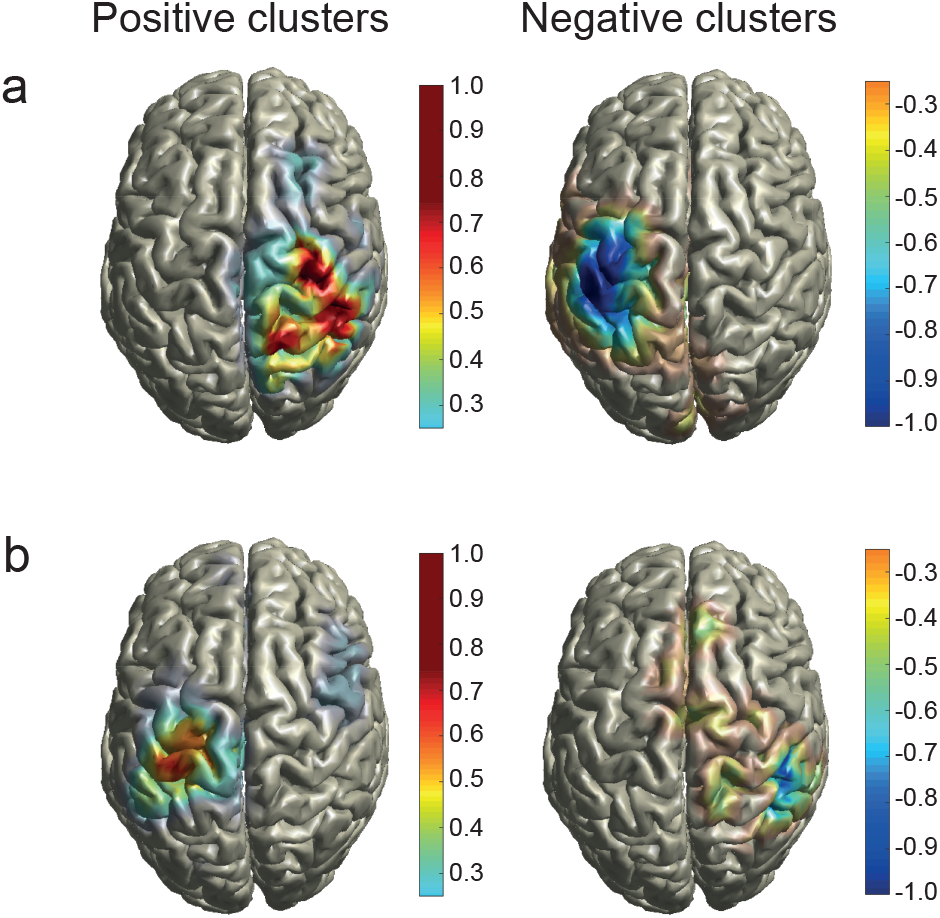
Clusters of Spearman’s rho averaged across conditions. Rho was computed by correlating source-level representation of ‘go’ ERF and RT (**Methods**). **a**) Clusters around 200 ms post-’go’ cue (see **Tbl. S3** for time spans). **b**) Clusters around 400 ms post-’go’ cue (see **Tbl. S3** for time spans). Colorbar to be interpreted as discrete as it depicts the number of conditions in which a source is part of a cluster: ± 0.25: one condition, ± 0.5: two conditions, ± 0.75: three conditions, ± 1: four conditions.

The early *positive* clusters cover parts of right motor cortex as well as parietal cortex (**Fig. 4a**, left). The location of these clusters is ipsilateral to the hand performing the button press which is in line with reports of ipsilateral motor-related activity observed at both the single-cell level^36^ and with MEG^37,38^, that may precede the onset of finger movements^39^. Our result likely is another manifestation of this phenomenon.

Note that the interpretation of these early clusters as parts of a single dipole, as may be inferred based on the sensor level analysis, is inadequate since the activities that underlie the positive and negative clusters originate from different hemispheres. **Fig. 5** depicts the location of all positive and negative early clusters in the four separate conditions. Although there are some differences in cluster extent between conditions, e.g. the positive cluster in auditory flipped exponential and the negative cluster in the auditory exponential condition both include parts of the frontal lobe, the averages across conditions in **Fig. 4** give a reasonable estimate of the early clusters’ location. [N.b. the (minor) extensions of the positive cluster to the left hemisphere in the visual exponential condition and of the negative cluster to the right hemisphere in the auditory exponential condition should be considered as artifacts due to leakage.]

**Fig. 5.**
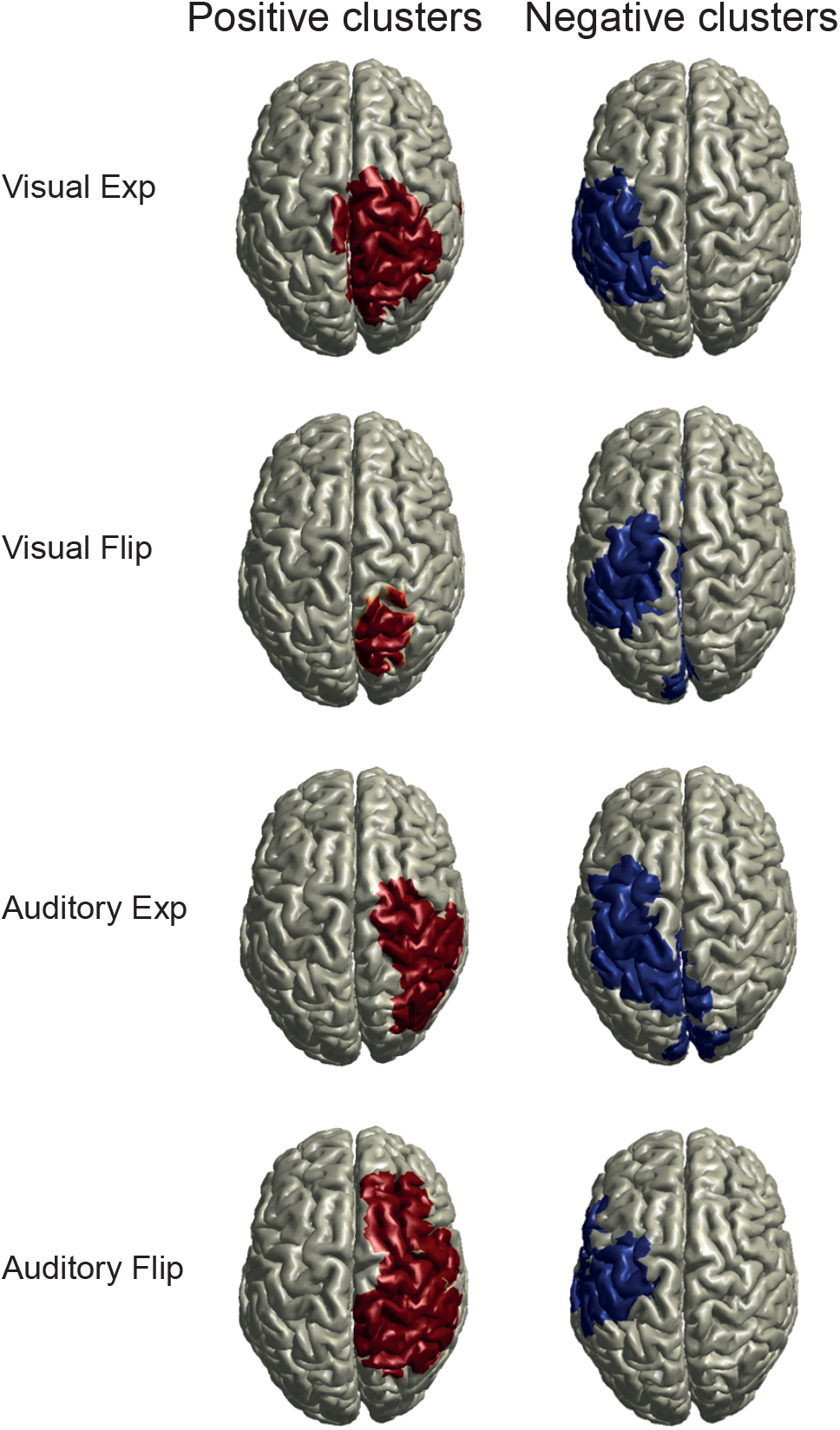
Clusters of Spearman’s rho around 200 ms post-’go’ cue. Rho was computed by correlating source-level representation of ‘go’ ERF and RT (**Methods**). See **Tbl. S3** for time spans of clusters.

The late clusters around 400 ms post-’go’ cue are depicted as across-condition averages in **Fig. 4b**. The late *positive* clusters are located over left primary motor and somatosensory cortex (**Fig. 4b**, left), whereas the late *negative* clusters are located over lateral posterior parietal cortex (**Fig. 4b**, right). The distribution of sources for both positive and negative late clusters is more variable across conditions (**Fig. S6**) than in the case of the early clusters (**Fig. 5**). E.g. in the auditory exponential condition, the positive cluster comprises of sources in left parietal cortex and also right latero-frontal cortex (as well as a small left centro-frontal component) (**Fig. S6**, left). Potential reasons for this variation in location within a single cluster are offered in a dedicated section below. Still, the locations of early and late clusters mostly overlap (compare **Fig. 4a**, left with 4**b**, right; and **Fig. 4a**, right with 4**b**, left). The overlap between *early* negative (**Fig. 4a**, right) and *late* positive (**Fig. 4b**, left) clusters supports the interpretation of these clusters as motor and somatosensory components of the button ERF.

The late negative clusters over right parietal cortex (**Fig. 4b**, right) are in agreement with the probability-sensitive ERP component P300. The P300 (P3b) originates from parietal areas^25,40^ and is commonly observed in the time span around 400 ms^19-21,23,31^. In contrast, the N2c component, which is also sensitive to probability^24^, is argued to originate from frontal cortex^41^ and peaks earlier than the P300, i.e. around 250 to 350 ms after a simple stimulus^42^. Work using EEG promotes a negative correlation between P300 amplitude and probability^19^. Here, using MEG, the negative correlation between the P300 and RT implies that the component’s amplitude positively covaries with probability density, i.e. the P300’s amplitude is large where expectancy is high and vice versa.

Note that as is the case in both early clusters, the average activity of positive and negative late clusters (**Fig. 4b**) originates (mostly, see **Fig. S6**) from different hemispheres.

As a sanity check of the above source-level analysis, Spearman’s rho was investigated. The distribution of mean rho averaged over the time spans of the identified clusters (**Tbl. S3**) was centered around zero (**Fig. S7**) without excessive skewing towards negative or positive values. The values of mean rho that comprised each cluster show a clear shift away from zero (**Fig. S8**), reflecting the sign of the identified clusters. The number of sources varies across clusters (compare e.g. positive cluster 1 between visual exponential and visual flipped exponential, **Fig. S8**) which is also reflected in the source projection plots (e.g. **Fig. 5** positive cluster visual exponential and flipped exponential). Despite these differences in source number, the clusters appear around the same cortical locations.

The correlation-based analysis in source-space identified a second positive cluster of rho (correlation between RT and ERF) around 200 ms post-’go’ cue (**Tbls. S3** and **S4**). This cluster comprised of sources in the cerebellum (**Fig. 6**). The consistency of this finding across all four conditions supports the functional role of the cerebellum in both, motor control^43^ and in the estimation of time^44^.

**Fig. 6.**
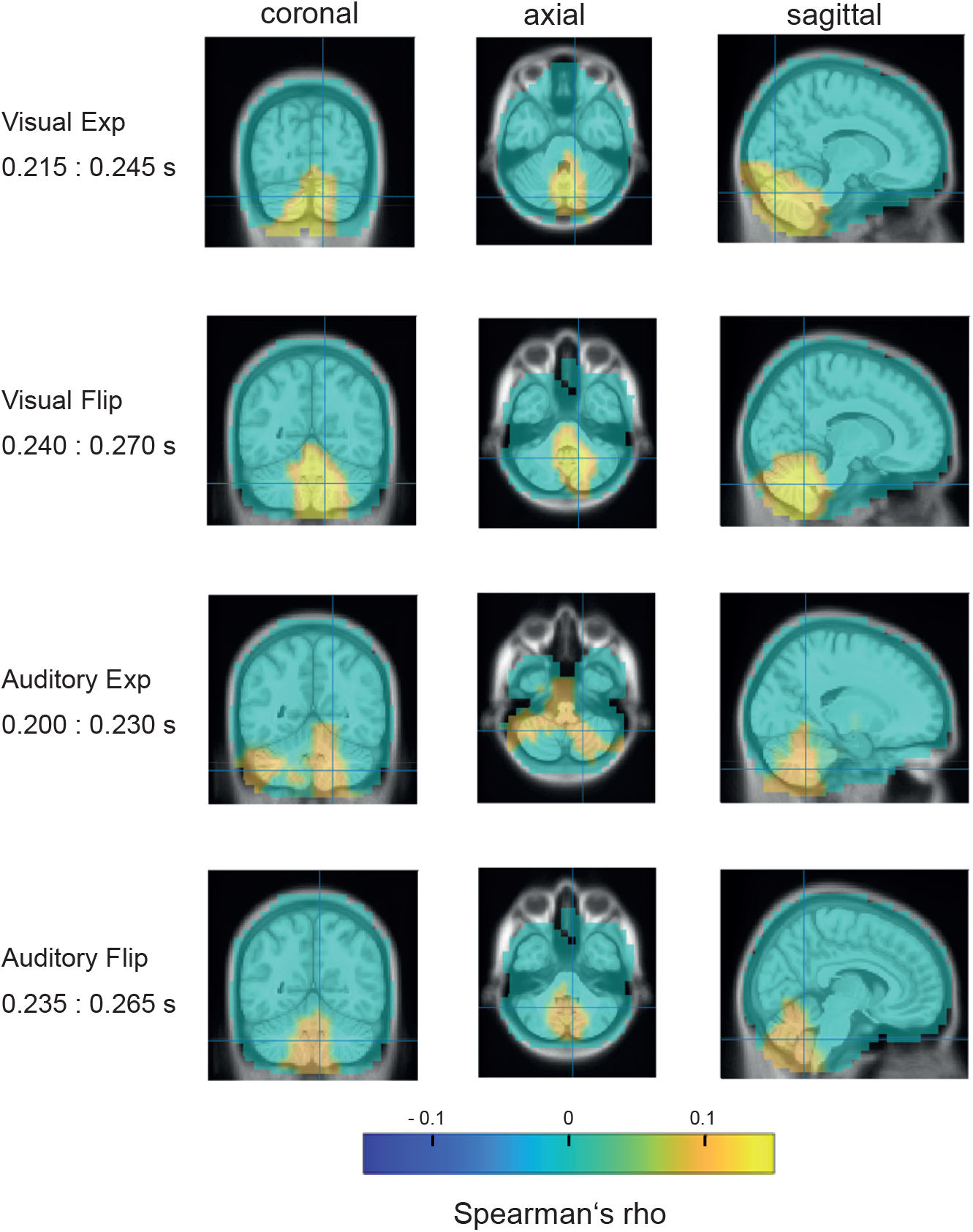
Cerebellar clusters of Spearman’s rho computed between ERF and RT around 200 ms post-’go’ cue (see **Tbl. S3** for time spans and **Methods** for details on correlation). Coronal: view from back, axial: view from top, sagittal: view from right.

In sum, the analysis of time-locked data identified significant correlations between the ERF and RT. There were clusters over left sensory-motor cortex and clusters over right parietal cortex. The clusters over left sensory-motor cortex, early negative and late positive, and the positive cluster located in the cerebellum reflect the sensitivity of RT to ‘go’ time probability. The finding of a late negative cluster in all conditions, irrespective of ‘go’ time distribution and sensory modality, consistently points to an involvement of right parietal cortex in temporal anticipation. No correlates between the ERF and RT were found in the time span before the ‘go’ cue, indicating absence of preparatory activity, such as the CNV, in time-locked data. Finally, no correlation was observed in early sensory ERF components (<150 ms) in visual or auditory areas, indicating that these sensory responses were not modulated by probability density.

### Correlation between 1/PDF and MEG data

The analysis of the evoked MEG data demonstrated that ERF components after the onset of the ‘go cue’ are significantly correlated with RT. As the initial behavioral analysis identified 1/PDF as an adequate explanatory variable of RT over ‘go’ time (**Fig. 1e** and **f**), the question naturally followed whether this simple linear model would also show similar correlations with the MEG data as with the RT.

The answer is not obvious for the following reason. In the correlation analysis of MEG with RT, Spearman’s rho was first computed at the single-subject level and then the statistics were performed across subjects. In the initial behavioral analysis, the 1/PDF model was fit to the group-level data. This means that the average RT curve for each subject was computed, then these curves were again averaged on the group-level and a single 1/PDF model was fit to them (**Methods**). This group-level model gave a very accurate fit with high R^2^ values. However, when this 1/PDF model was fit on the single-subject level, although it qualitatively followed the individual RT curve, quantitatively it had much smaller R^2^ values due to the higher levels of variability and noise (**Fig. S9**). So the question can be reformulated into whether this higher variability and noise at the single-subject level allows the 1/PDF models to have the quantitative accuracy required to capture the correlations with the MEG data that were observed in the RT-MEG correlation analysis (see **Suppl. Methods** for further details).

### Event-related field components – correlation with 1/PDF at the sensor level

In this section we employ the same correlation-based approach but using the 1/PDF as a regressor instead of RT. First, the single-subject RT curves were fit with the reciprocal probabilistically blurred PDF. The 1/PDF models and the MEG data time-locked to the ‘go’ cue (and baselined using a pre-’go’ baseline) were aggregated in 30 frames (see above, **Methods**). For each channel-time duplet, Spearman’s rho was computed between the aggregated model values and the aggregated MEG data followed by a cluster-based permutation test on rho. Similar to the RT correlation case, a pattern of early (∼ 200 ms) and late (∼ 400 ms) positive clusters (**Fig. S10a**) and negative clusters (**Fig. S10b**) was identified (**Tbl. S6**) with topographies (**Fig. S10c** and **d** and **Fig. S11a** and **b**) similar to those in the RT case (**Figs. 3** and **S2**). These similarities between the two regressors’ clusters provides first evidence that the 1/PDF model is not only a valid model of RT behavior but it also points to the neural processes involved in the generation of reaction times. Plots of rho and the ERF over time, both averaged across each clusters’ MEG channels, further support this finding (**Fig. S12**). Rho averaged within cluster channels has a similar time course in both RT and 1/PDF regressor cases (compare **Fig. S12a**, left column, with **Fig. S12b**, left column). Likewise, the ERF averaged within cluster channels is similar for both regressors, reflecting that both regressors identified mostly the same cluster channels (compare **Fig. S12a**, right column, with **Fig. S12b**, right column). Taken together, the similarities in RT-based and 1/PDF-based correlation results lend support to the interchangeability of the two regressors.

### Event-related field components – correlation with 1/PDF at the source level

Next, the above cluster-based analysis was performed on Spearman’s rho computed by correlating the fitted 1/PDF model and the source-space representation of the ERF (**Fig. S13, Methods**). The cluster-based permutation test on rho identified a pattern of clusters around 200 ms (**Tbls. S7** and **S8**) that is highly similar to the one observed in the above analysis of rho computed on RT as a regressor (**Tbl. 1**): the sources that comprise the positive clusters are located over right parietal cortex; the sources that comprise the negative clusters align well with left parietal cortex including left pre-motor and primary motor cortex (**Fig. 7**).

**Fig. 7.**
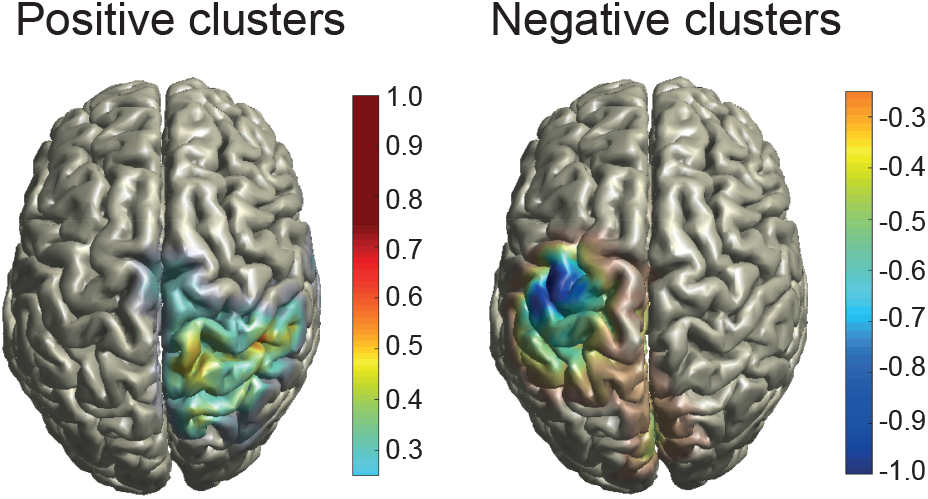
Clusters of Spearman’s rho averaged across conditions. Rho was computed by correlating source-level representation of ERF and fit 1/PDF (**Methods**). **a**) Clusters around 200 ms post-‘go’ cue (see **Tbl. S7** for time spans). **b**) Clusters around 200 ms post-’go’ cue (see **Tbl. S7** for time spans). Colorbar to be interpreted as discrete as it depicts the number of conditions in which a source is part of a cluster: ± 0.25: one condition, ± 0.5: two conditions, ± 0.75: three conditions, ± 1: four conditions.

As was to be expected, given that the 1/PDF regressor is an adequate model of RT, in all four conditions the locations of positive and negative clusters in the 1/PDF regressor case (**Fig. 8**) closely resembled those in the RT regressor case (**Fig. 5**). As a sanity check, histograms of Spearman’s rho (pooled across sources within each cluster’s time span) were investigated (**Fig.S14**). They depict a close-to-symmetric distribution of rho centered around zero in all cases. Histograms of all rho values per cluster (**Fig. S15**) show a shift in average rho away from zero as well as a difference in the number of sources between the clusters. All of these findings closely resemble those from the time-locked analysis using RT as a regressor (see above).

**Fig. 8.**
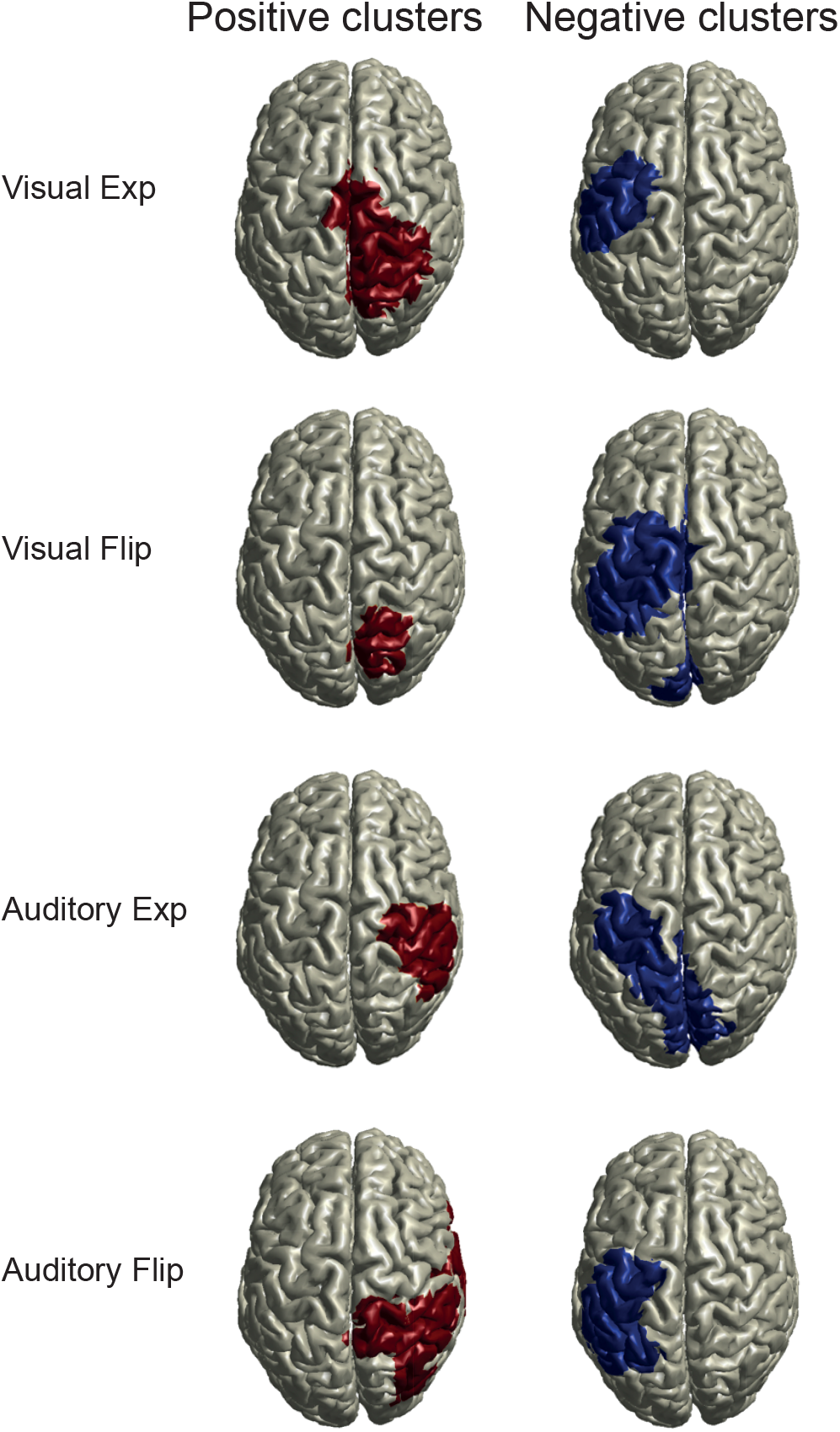
Clusters of Spearman’s rho around 200 ms post-’go’ cue. Rho was computed by correlating source-level representation of ‘go’ ERF and fit 1/PDF (**Methods**). See **Tbl. S7** for time spans of clusters.

## Discussion

We investigated the neural correlates of temporal anticipation. Reaction times to visual and auditory cues followed the reciprocal event probability density. The temporal dynamics in right parietal cortex correlated with RT to anticipated events. More specifically, time-locked activity in right parietal cortex after the event occurred was correlated with probability density, hinting at a functional role of late ERP component P300 in temporal expectancy. This result is independent of sensory modality, suggesting a central role of parietal cortex in the estimation of probability density.

### Reciprocal PDF but not HR captures reaction times

We replicated our previous result that RT modulation is captured by a model based on the 1/PDF, whereas the canonical HR-based model failed to fit RT. Although the HR model makes intuitive sense, its failure at the behavioral level renders it an inadequate model of event anticipation. The 1/PDF model is based on the event PDF itself, a variable that can be built on computational primitives such as the counting of events over time. This computation can easily be implemented in an elementary neural network^45^ indicating computational parsimony and biological plausibility. We used the 1/PDF model as a regressor on neural data.

### No effect of probability density on pre-stimulus time-locked neural activity

Our analysis did not identify a significant correlation between pre-stimulus event-related components, such as the contingent negative variation (CNV) and the lateralized readiness potential (LRP), with RT or its 1/PDF model. In the case of the LRP this is not surprising, as the LRP is commonly explored using contralateral-ipsilateral response difference waves in both EEG^27^ and MEG^46^ work to avoid confounding activity introduced by the stimulus. In the present experiment, participants responded only with one hand and the subtractive analysis approach was not possible. In previous work on the CNV, its sensitivity to event probability was shown for between-stimuli time spans larger than 1 s^6,47^. In fact, when stimuli are separated by several seconds, the CNV may consist of two components, an early one that is associated with processing of the first stimulus and a second one that precedes the second stimulus and is associated with response preparation^29^. Our task employed the ‘go’ time span of 0.4 to 1.4 s which may be too short for a probability-driven modulation of the CNV wave to build up. In addition, our design employed full distributions defined over 60 ‘go’ times (∆ t = 1/60 s) whereas commonly only a small number of different ‘go’ times are presented^6,47^. The modulation of response preparation, e.g. by attribution of attention to only a few time points, may likely differ from its (quasi-)continuous counterpart investigated here, resulting in the absence of significant CNV modulation by event probability density.

### No effect of probability density on early ERF components

We next investigated whether early post-stimulus ERF components are modulated by event probability density. This hypothesis was motivated based on work proposing early cortical processing of event probability at the single-cell^1,4^ and population response^5^ levels, and specifically the sensitivity to attention of e.g. the visual P1^23,48-50^ and the auditory N1^51^ wave. We observed no significant correlation between midlatency (10 to 50 ms) or longlatency (50 to ∼150 ms) ERF components and probability density. The common finding of P1 amplitude increase is associated with enhanced stimulus processing and is thought to be driven by spatial^48^ and spatial-temporal^23,50^ attention. In the temporal attention case, in Rohenkohl and Nobre’s work, the event occurred at one of only two different stimulus time points and did not require complex probability estimation^23^, whereas our task required the distribution of attention over the time span of one second based on the participant’s estimate of event probability. Our stimuli are determined by event probability density functions and may therefore be an ecologically more realistic approximation to environmental dynamics^52^. However, the temporal-probabilistic structure embedded in our task may have been too complex to evoke these ERP effects. As Luck et al. put it: “…these early ERP components are small and may be influenced by attention only when attention is very highly focused”^53^. If this is indeed the case, then, given the obvious modulation of RT, the absence of early ERP modulation suggests that either a) the effect may still be there but our design is not sensitive enough to pick it up, or, b) the effect is not there because enhanced stimulus processing at very early cortical levels is not crucial in the task. The later interpretation may be more adequate since Rohenkohl and Nobre investigated perceptual decision making (discrimination of visual stimuli)^23^ whereas our simple RT task contains no choice component. Thus, it may be that in temporal-probabilistic inference, the brain does not benefit from adaptive stimulus-related processing in early cortical areas. A more straightforward reason for the absence of effect may be that these early components decrease in amplitude as the interval between two stimuli decreases^25^ which may interfere with putative effects of probability density. In sum, the analysis of time-locked data did not produce evidence for early cortical stimulus processing that is driven by event probability density.

### Probability density modulates parietal P300 wave

Our analysis identified a modulation of the P300 (P3b) ERF component by event probability density. The fact that the P300 occurred *after* the response was given raises the question of the component’s functional role in temporal anticipation. In more complex tasks from decision-making involving choice, where the P300 occurs *before* the choice is instantiated, the component is argued to (partly) represent decision processes^54^. Our simple task only required a decision regarding the question: “Shall I press the button now?” which is solely contingent on the occurrence of the ‘go’ cue. We therefore favor other common interpretations such as that the P300 may represent processes involved in the updating of working memory^55-57^ or information updating in general. In this regard the concept of context updating^58^ proposes an attention-driven comparison between current sensory input and the representation of past sensory events. This hypothetical comparison-and-updating process could be conceptualized in our task by a trial-by-trial process model that updates probability estimates over time. This process model could possibly predict the trial-by-trial P300. Yet it would remain unclear how these hypothetical processes link causally to the observed effects on the P300. To better understand the functional role of this prominent component in temporal anticipation, a targeted EEG experiment could be performed in which the presumed updating processes reflected in the P300 are perturbed using transcranial magnetic stimulation (TMS). Nonetheless, the finding that the P300 modulation by probability density is very similar in vision and audition is in line with this late component’s known independence from input modality^25^ which suggests high-level processing.

We further identified an interesting anatomical relationship between early and late clusters. The *early* positive clusters cover parts of parietal cortex, as well as motor cortex (**Fig. 4a**, left). The location of these clusters is ipsilateral to the hand performing the button press. The ipsilateral activity prior to a motor action has been proposed to represent suppression of contralateral movement^59^, but its precise functional significance remains unclear. Given the observed modulation of the P300 by probability density, an alternative view would be that the parietal components of these early positive clusters do not reflect movement suppression but are functionally related to the presumed information updating processes reflected in the late negative clusters, i.e. the P300.

In summary, the time-locked analysis revealed that the P300 co-varies with expectancy and that it originates from the right parietal lobe associating this cortical area with the later processing stages of event probability density.

### HR and 1/PDF as regressors in psychometric-neurometric mapping

The HR has been proposed as a model of anticipatory neural activity at the single-cell and population response level and at time spans ranging from hundreds of milliseconds to multiple seconds. Although comparison of results across different levels of analysis and across different time spans requires caution, the obvious difference between work promoting the HR and the present work promoting the 1/PDF variable is this choice of regressor. The HR variable and the 1/PDF variable differ in the ‘go’ time distributions used here (exponential case: U-shaped vs. quasi-monotonically increasing over time, flipped exponential case: convex vs. concave shape, **Fig. 1**). These qualitative differences between HR and 1/PDF exist for several distributions commonly used in experiments on temporal anticipation, such as the Gaussian^15,16^ and qualitatively similar distributions^1^, and also the Weibull^2,5^ and uniform^15^ distributions. These differences intuitively highlight the dependence of psychometric-neurometric mapping approaches on the choice of regressor. Note that we do not imply here that the choice of regressor renders the findings of previous work questionable per se. On the contrary, we suggest that the neural activity identified by the use of a HR-based regressor may in some cases be explained by other, simpler variables. In the case of event certainty, for many distributions (e.g. Gaussian and uniform), the HR variable monotonically increases over time. Correlation between HR and neural data may therefore identify a ramping activity that may or may not be driven by HR. On a related note, work on anticipation is commonly limited to only a single sensory modality. Again, this may mislead inference since it is not readily apparent how modality-specific contingencies can be separated from activity related to the processing of event probability density. Furthermore, the impact of probability on neural activity is sometimes investigated in discrete settings^6,10,60^ where HR is computed over discrete time^6,10^. Given the brain’s immense capacity for accurate processing of time-based information^61-63^, it is not obvious why a concept based on continuous time (the HR) should per se be adequate as a model of events that are highly discretized in time. In sum, common psychometric-neurometric mapping strategies critically depend on the choice of regressor. In our results, the good performance of the 1/PDF as both a model of average RT and as a regressor on averaged ERF data was contrasted by the less adequate fit of 1/PDF to single-trial RT data. This limitation of our descriptive model again illustrates the challenges of regressor choice. The aim of future work will be the development of a PDF-based model that gives a per-trial prediction of RT. Taken together, our results are more in line with work suggesting that probability density itself is reflected in neural anticipatory activity^64,65^, but not HR.

In conclusion, we report a supra-modal neural representation of event probability density in right parietal cortex. This result is supported by the modulation of ERP component P300. Overall, this work contributes to the understanding of the cortical processes involved in temporal anticipation.

## Materials and Methods

The experiments were approved by the Ethics Committee of the University Hospital Frankfurt. Written informed consent was given by all participants prior to the experiment.

### Participants

24 healthy adults (15 female), aged 21-34 years, mean age 27 years, participated in the experiment. All were right-handed and had normal or corrected-to-normal vision, reported no hearing impairment and no history of neurological disorder. Participants received € 15 per hour. One subject was excluded from the source-level analysis of MEG data because the anatomical MRI data were corrupted.

### Experimental task and stimuli

In the MEG booth, participants performed visual and auditory blocks of trials of a ‘set’ - ‘go’ task. In the task, a ‘set’ cue was followed by a ‘go’ cue (**Fig. 1a**). Participants were asked to respond as quickly as possible with a button press (Current Designs Inc., Philadelphia, PA, USA) to the ‘go’ cue using their right index finger. Participants were instructed to foveate a central black fixation dot and restrict blinking to the timespan immediately following a button press. In 10 % of trials, no ‘go’ cue was presented. In these catch trials, participants were asked to not press a button. This small percentage of catch trials was added to avoid possible strong effects of event certainty towards the end of the ‘go’ time span^15^. A small black circle around the central fixation dot was presented onscreen for 200 ms after a button press indicating the end of the trial. In the case of a catch trial, the circle appeared after the longest possible ‘go’ time, i.e. 1.4 s. The intertrial interval (ITI) was defined by the onset of the small black circle and the ‘set’ cue of the following trial. The ITI was drawn randomly from a uniform distribution (range 1.4 to 2.4. s, discretized in steps of 200 ms).

### Visual stimuli

Two simultaneously presented checker boards served as both ‘set’ and ‘go’ cues. They were projected (refresh rate 60 Hz) to the back of a gray semi-translucent screen located at a fixed distance of approximately 53 cm from the participants’ eyes. Each cue was onscreen for 50 ms. The checker boards subtended approximately 6.5 × 6.5° visual angle and comprised of 5 × 5 small black and white squares of equal size. The center of the checker boards was located to the left and right of a central fixation dot at a horizonal distance of approximately 7° visual angle and a vertical distance of 0° visual angle. The black and white pattern of the checker boards was inverted between ‘set’ and ‘go’ cues.

### Auditory stimuli

White noise bursts of 50 ms length served as ‘set’ and ‘go’ cues. Each burst featured an 8 ms cosine ramp at beginning and end. The bursts were presented at approx. 60 dB SPL above hearing threshold as determined by pure tone audiometry (1 kHz, staircase procedure). All auditory stimuli were output via a RME Fireface UCX interface to a headphone amp (Lake People GT-109) and delivered diotically via a MEG-compatible tube-based system (Eartone Gold 3A 3C, Etymotic Research, Elk Grove Village, IL, USA). Visual and auditory stimuli were generated using MatLab (The MathWorks, Natick, MA, USA) and the Psychophysics Toolbox^66^ on a Fujitsy Celsius R940 computer running Windows 7 (64 bit).

### Temporal probabilities

The time between ‘set’ and ‘go’ cues, the ‘go’ time, was a random variable, drawn from either an exponential distribution (Equation 1) with parameter *l* = 0.33 or from its left-right flipped counterpart.

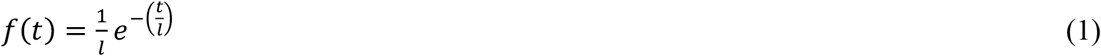

Both distributions were delayed by 0.4 s resulting in a range of ‘go’ times from 0.4 s to 1.4 s. Sequential effects were reduced by the constraint that no more than two consecutive trials were allowed the same ‘go’ time. The ‘go’ time distribution was fixed for a pair of consecutive blocks of trials. Per participant the experiment consisted of four visual and four auditory blocks. A single block consisted of 200 trials of which 20 did not feature a ‘go’ cue (catch trials). Per sensory modality, in two blocks of trials, the ‘go’ times were randomly drawn from the exponential distribution and in the other two blocks they were randomly drawn from the flipped-exponential distribution as described above.

### Models of reaction time

Hazard-rate-based and PDF-based models of RT were constructed to investigate the effect of the ‘go’ time distribution on event anticipation. The presented exponential and flipped exponential ‘go’ time distributions are characterized by three functions, the probability density function (PDF), the cumulative distribution function (CDF), and the hazard rate (HR):

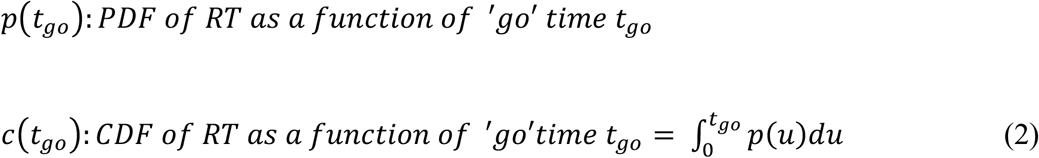

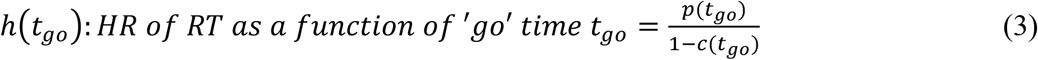

### Mirrored temporally blurred hazard rate

To arrive at the temporally blurred HR, each ‘go’ time PDF was blurred by a Gaussian uncertainty kernel whose standard deviation linearly increases with elapsed time from a reference time point: σ = φ ∙ *t*. Here, *t* is the elapsed time and φ is a scale factor by which the standard deviation σ of the Gaussian kernel increases. The equations for the corresponding temporally blurred functions are:

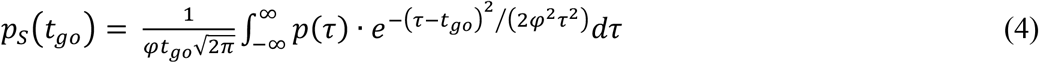

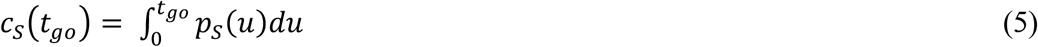

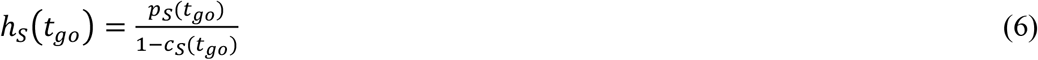

For a given ‘go’ time *t*_*go*_ the PDF is convolved with a Gaussian kernel centered at *t*_*go*_ (Equation 4). At *t*_*go*_ = 0.4 *s* after ‘set’ cue onset the kernel has standard deviation *φ* ∙ 0.4. Similarly at *t*_*go*_ = 1.4 *s* the kernel has standard deviation φ ∙ 1.4. In the computation of the subjective PDF, the definition of the PDF was extended to the left and right of the ‘go’ time range:

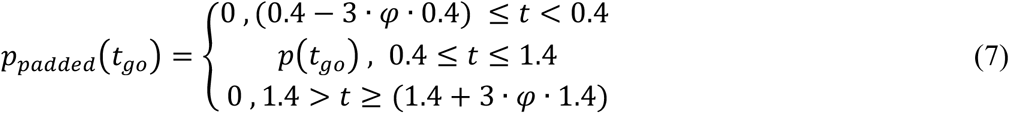

The extensions were equal to three standard deviations of the Gaussian kernel (encapsulating 99.7 % of the Gaussian uncertainty function) at the shortest and longest ‘go’ times. Then the integral in Equation (5) was computed between these new extrema [(0.4 − 3 ∙ *φ* ∙ 0.4), (1.4 + 3 ∙ *φ* ∙ 1.4)] instead of the impractical interval of minus to plus infinity. The selection of *φ* = 0.21 was consistent with previous research^2,16,67,68^. For *φ* = 0.21 the temporal range of the extended PDF (7) becomes [0.148, 2.28] s which is also the range of integration in the computation of the subjective PDF in Equation (4). The PDF of each distribution, *p*-*t*_*go*_. was normalized so that its integral from 0.4 s to 1.4 s was 0.9 which reflecte the 10% catch trials that did not feature a ‘go’ cue. The HR was computed based on the PDF and CDF (Equation 6). To arrive at the to-be-fit HR variable, the HR was mirrored around its mean (Equation 8).

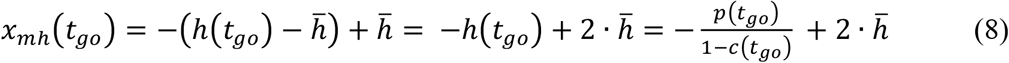

where

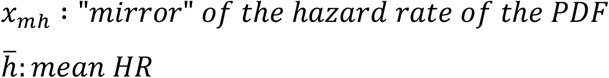

### Reciprocal probabilistically blurred event probability density function

Probabilistic blurring constitutes an alternative hypothesis to the temporal blurring described above. In probabilistic blurring, the uncertainty in elapsed time estimation depends on the probability density function of event occurrence: ‘go’ times with high probability of event occurrence are associated with low uncertainty in time estimation and vice versa, irrespective of the ‘go’ time duration^16^. In probabilistic blurring, the standard deviation of the Gaussian kernel scales according to the PDF of event occurrence. In order to use realistic values for the standard deviation of the blurring Gaussian kernel, the minimum and maximum values were set accordingly to the temporal blurring case as

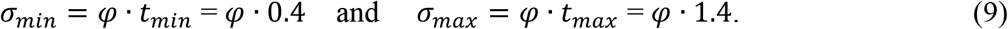

The value of φ was likewise set to 0.21. The PDF under investigation was then scaled so that its minimum value is σ_*min*_ and its maximum value σ_*max*_

If *pmin* and *p*_*max*_ are the minimum and maximum values respectively of the PDF under investigation then the function used for computing the standard deviation of the Gaussian kernel based on the PDF *p*(*t*) was defined as:

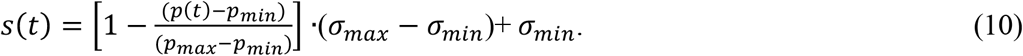

The term inside the brackets demonstrates that when the probability *p*(*t*) is low the standard deviation of the Gaussian kernel approaches σ_*max*_ while when the probability increases, *s*(*t*) approaches σ_*min*_.

Based on Equation (10) for determining the standard deviation of the Gaussian kernel the probabilistically blurred PDF *p*_*p*_(*t*) was computed as:

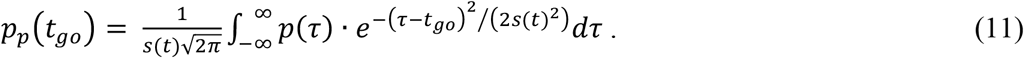

Finally, in order to implement the Gaussian blurring of Equation (11) at the extrema of ‘go’-times, the definition of the PDF was extended to the left and right of the actual stimulus presentation interval by three standard deviations of the corresponding smoothing Gaussian kernels, similar to the temporally blurred case described in Equation (7) as:

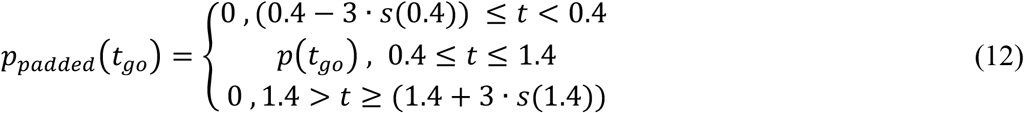

The extensions of the ‘go’ times range depend on the standard deviation function *s*(*t*), which itself depends on the probability density function. To arrive at the to-be-fit PDF variable, the reciprocal of the PDF was computed: 1 / PDF (Equation 13).

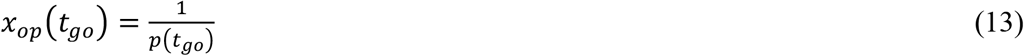

Where

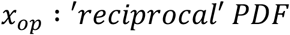

### Modeling RT with a linear model

Both variables, the mirrored, temporally blurred HR and the probabilistically blurred 1/PDF were fit to RT data (first aggregated within ‘go’ times within participants, then averaged across participants) using a linear model. An Ordinary Least Squares (OLS) regression was employed for the computation of the regression coefficients using the MatLab (The MathWorks, Natick MA, USA) *fit* function. Adjusted *R*^1^ was used as a measure of goodness-of-fit for comparing the models’ relation to RT.

### MEG data acquisition

Neuromagnetic activity was recorded with a 275-channel system (VSM MedTech Omega, Coquitlam, Canada) equipped with axial gradiometers distributed in a helmet across the scalp in a magnetically shielded room at the Brain Imaging Center, Frankfurt. MEG data were recorded continuously with a sampling rate of 1,200 Hz. Participants were seated in an upright position and were asked to remain still during blocks of trials. Head position relative to the MEG sensors was controlled and continually monitored during each experimental block using three position indicator coils in the anatomical fiducial locations (left and right pre-auricular points and nasion). Head position was corrected if necessary between blocks using the fieldtrip toolbox^35^. Electro-cardiogram and vertical and horizontal electro-oculogram were also measured at 1,200 Hz to identify eye blinks and movements, and the heartbeat in analysis.

### MEG data preprocessing

Continuous data were down-sampled to 600 Hz. Data were epoched separately with respect to ‘set’ and ‘go’ cues and to the button press (-0.5 to 0.72 s). Artifactual epoches and noisy MEG channels were rejected based on visual data inspection using Fieldtrip’s visual artifact rejection routines. Based on visual inspection, trials and MEG channels that featured periods of high variance due to e.g. eyeblinks or excessive movement, during the time span of interest were discarded from the analysis. Heartbeat artifacts were removed using independent component analysis. Drifts in MEG channels were eliminated by high-pass filtering at 0.2 Hz. Muscle activity was eliminated by low-pass filtering at 110 Hz. Finally, only trials with 0.05 s < RT < 0.75 s were used in the analysis of MEG data and in modeling of RT. The selection process resulted in the removal of N = 1204 trials, leaving N = 33354 trials for analysis.

### Magnetic resonance imaging data acquisition

To build participant-specific forward models for MEG source reconstruction, structural magnetic resonance imaging (MRI) scans (T1-weighted) were obtained for all 24 participants at the Brain Imaging Center, Frankfurt. MRI data were acquired on a Siemens 3T TRIO scanner with a voxel resolution of 1 × 1 × 1 mm^3^ on a 176 × 256 × 256 grid. Vitamin-E capsules were used to identify anatomic landmarks (left and right peri-auricular points and nasion).

### Event related fields analysis on sensor-space data

MEG data time-locked to the ‘go’ cue was aggregated within adjacent pairs of consecutive ‘go’ times (frames). For the first frame, the activity from all trials with ‘go’ times = [0.4, 0.4167] s were averaged over time; the second frame comprised ‘go’ times = [0.4333, 0.45] s and so forth. This procedure was also applied to the single-participant RT and to the single-participant fits of the 1/PDF model of RT. The aggregation reduced the number of unique ‘go’ times from 60 to 30 (= 30 frames). At the single-participant level, for each channel-by-time-point duplet, Spearman’s rank correlations were computed between the 30 frames of MEG data and the 30 averaged RTs (and the 1/PDF fitted to single-subject RT). Note that the resultant rho has the same dimensionality as the within-participant grand average of MEG data (channels-by-time-points). On all participants’ rho, statistical significance was tested using a cluster-based permutation test was run to identify channel-by-time clusters in which rho differs from zero (cluster-forming α = 0.05, 1000 permutations). Before the averaging of the ERF, the data was baselined either to the pre-SET or to the pre-GO period. The pre-SET period (t = [-0.5, 0] s) was selected in the analysis of activity preceding the ‘go’ cue, whereas the pre-GO period (t = [-0.5, 0] s) was selected in the analysis of activity following the ‘go’ cue.

### Event related fields analysis on source-space data

We used minimum variance beamformer LCMV^69^. The co-variance matrix used for computation of the spatial filter was derived from the average across trials. The time period used for computation of the covariance matrix was t = [-0.5, 0.9] s relative to the ‘go cue. The average, baselined ERF for the computation of the covariance matrix was derived by averaging all trials from all frames so that the spatial filters are common for all frames. Then the sensor-level average ERF for each frame was projected to source space through the common spatial filters. For each source location, Spearman correlation was computed between the average ERF across all frames and the corresponding RTs. This was repeated for each subject. Then the non-parametric cluster-based permutation statistics were computed to identify clusters of sources in which rho differs from zero (cluster-forming α = 0.05, 1000 permutations).

