## Supplementary Material for "Neural Correlates of Temporal Anticipation"

Matthias Grabenhorst[1,2] & Georgios Michalareas[2]

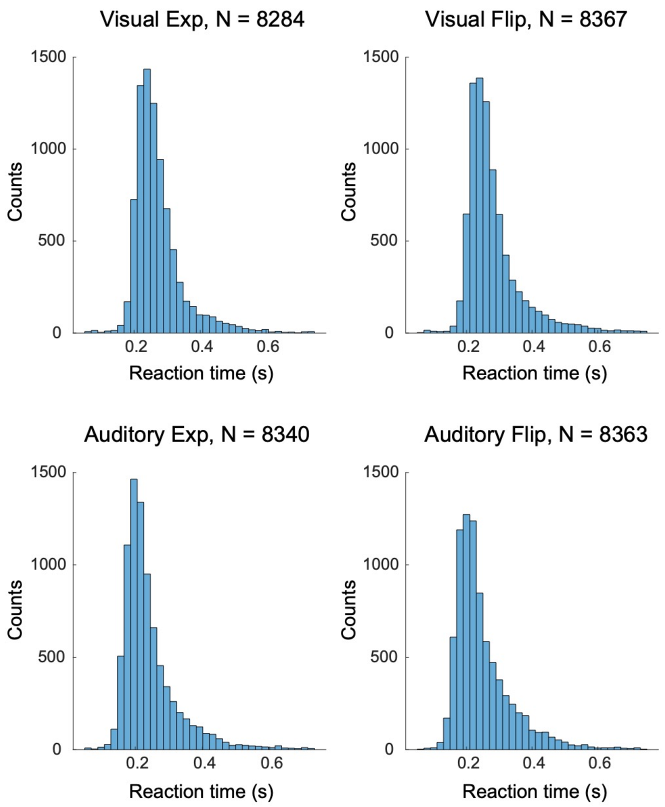


**Fig. S1** Histogram of reaction times (RT) for each experimental condition.

**Table S1** Cluster-based permutation test on Spearman's rho (correlation between sensor-level ERF of the 'go' cue and RT). ERF data baselined using pre-'go' cue baseline (**Methods**).

| Condition | Cue | Regressor | Positive cluster (*P*- value) | | Negative cluster (*P*- value) | |
|---|---|---|---|---|---|---|
| | | | #1 | #2 | #1 | #2 |
| Vis exp | 'go' | RT | 0.002 | 0.002 | 0.002 | (0.19) |
| Vis flip | 'go' | RT | 0.002 | 0.012 | 0.002 | 0.006 |
| Aud exp | 'go' | RT | 0.002 | 0.002 | 0.002 | 0.018 |
| Aud flip | 'go' | RT | 0.002 | 0.010 | 0.002 | --- |

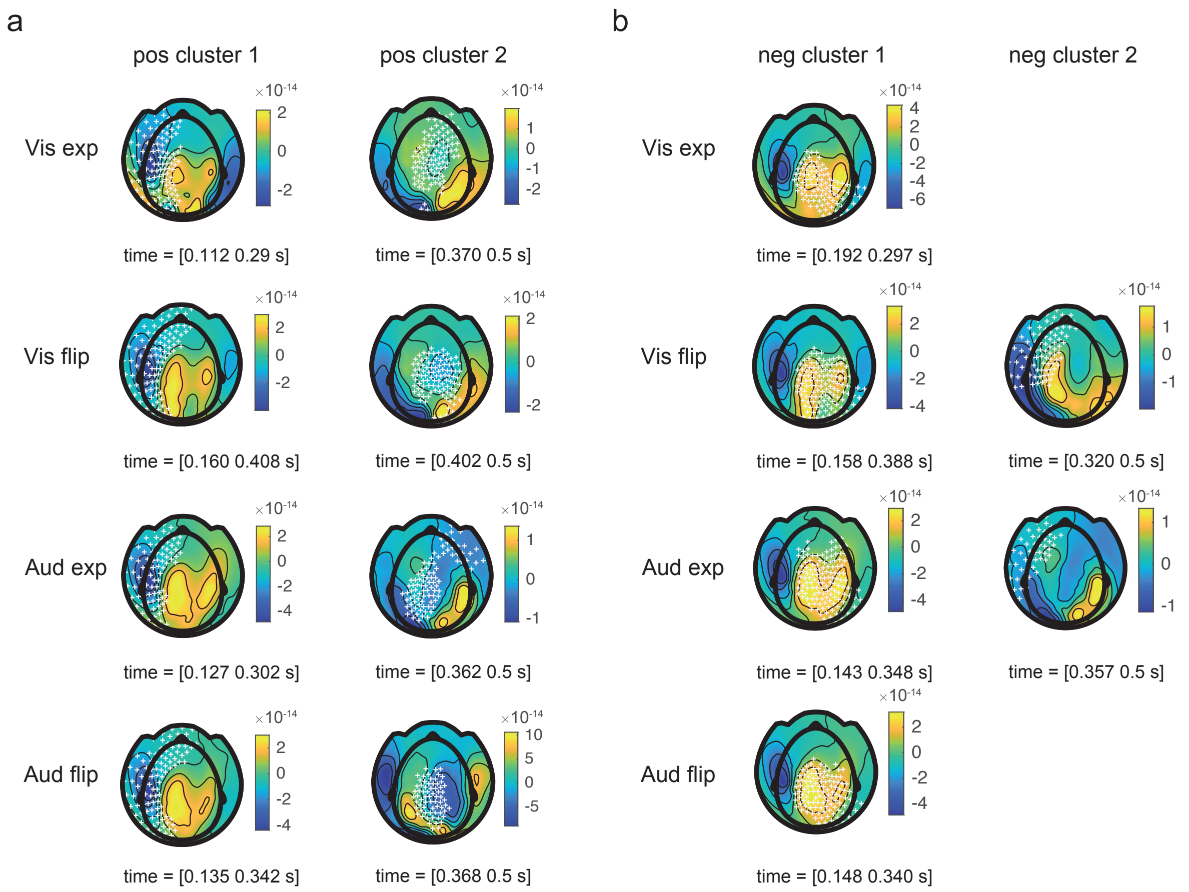


**Fig. S2** Cluster-based permutation test on Spearman's rho (correlation between ERF of the 'go' cue and RT). **a**) Topography plots of 'go' ERF, channels of positive cluster highlighted. **b**) Topography plots of 'go' ERF, channels of negative cluster highlighted. A minimum time span of 20 ms was set for a channel to be included in cluster. ERF data baselined using pre-'go' cue baseline (**Methods**).

**Table S2** Cluster-based permutation test on Spearman's rho. Correlation between sensor-level ERF of the 'go' cue and RT. ERF data baselined using pre-'set' cue baseline (**Methods**).

| Condition | Cue | Regressor | Positive cluster (*P*- value) | | Negative cluster (*P*- value) | |
|---|---|---|---|---|---|---|
| | | | #1 | #2 | #1 | #2 |
| Vis exp | 'go' | RT | 0.006 | 0.006 | 0.002 | (0.04) |
| Vis flip | 'go' | RT | 0.002 | 0.008 | 0.002 | 0.002 |
| Aud exp | 'go' | RT | 0.002 | 0.002 | 0.002 | --- |
| Aud flip | 'go' | RT | 0.002 | 0.002 | 0.002 | --- |

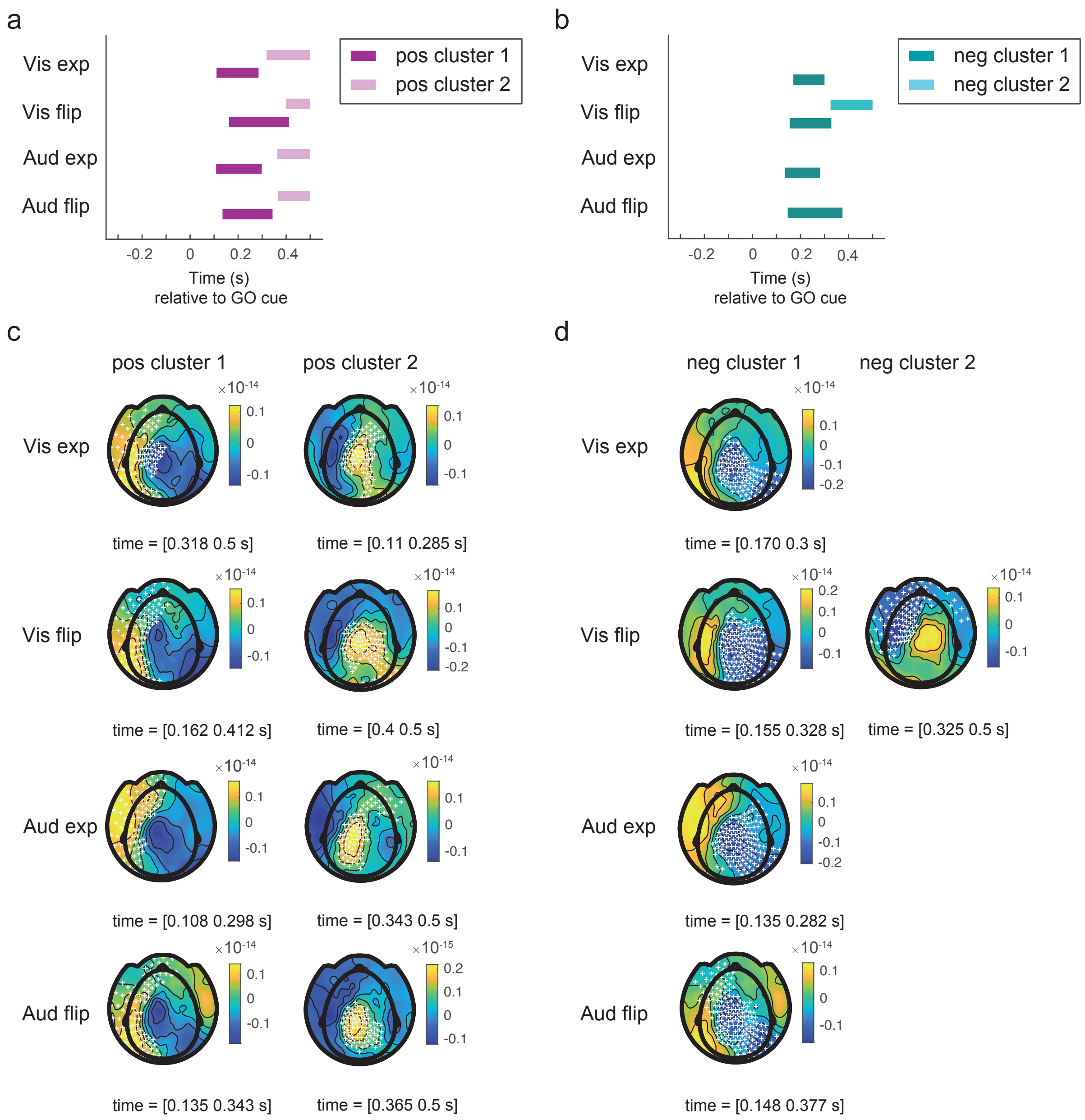


**Fig. S3** Cluster-based permutation test on Spearman's rho (correlation between ERF of the 'go' cue and RT). **a**) Positive clusters **b**) Negative clusters **c**) Topography plots of rho, channels of positive cluster highlighted **d**) Topography plots of rho, channels of negative cluster highlighted**.** A minimum time span of 20 ms was set for a channel to be included in cluster. ERF data baselined using pre-'set' cue baseline (**Methods**).

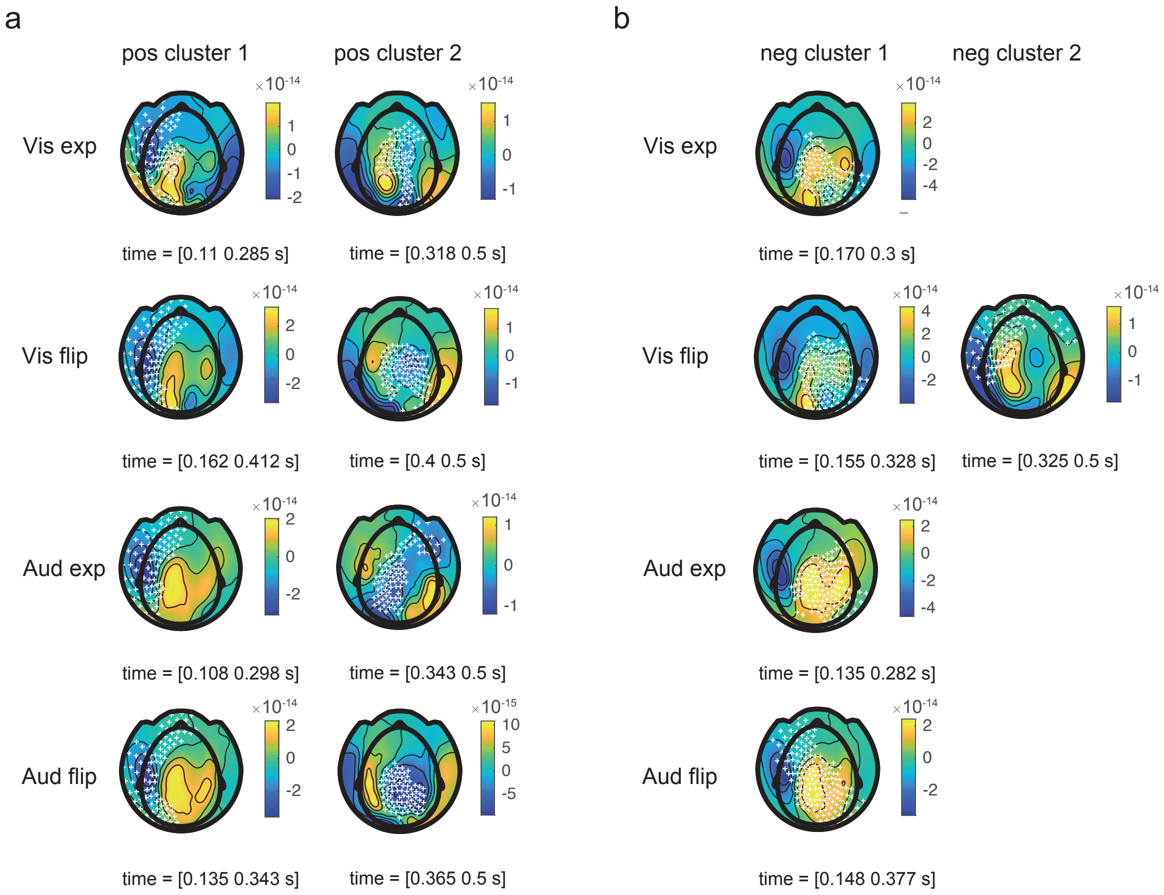


**Fig. S4** Cluster-based permutation test on Spearman's rho (correlation between ERF of the 'go' cue and fit RT). **a**) Topography plots of 'go' ERF, channels of positive cluster highlighted. **b**) Topography plots of 'go' ERF, channels of negative cluster highlighted. A minimum time span of 20 ms was set for a channel to be included in cluster. ERF data baselined using pre-'set' cue baseline (**Methods**).

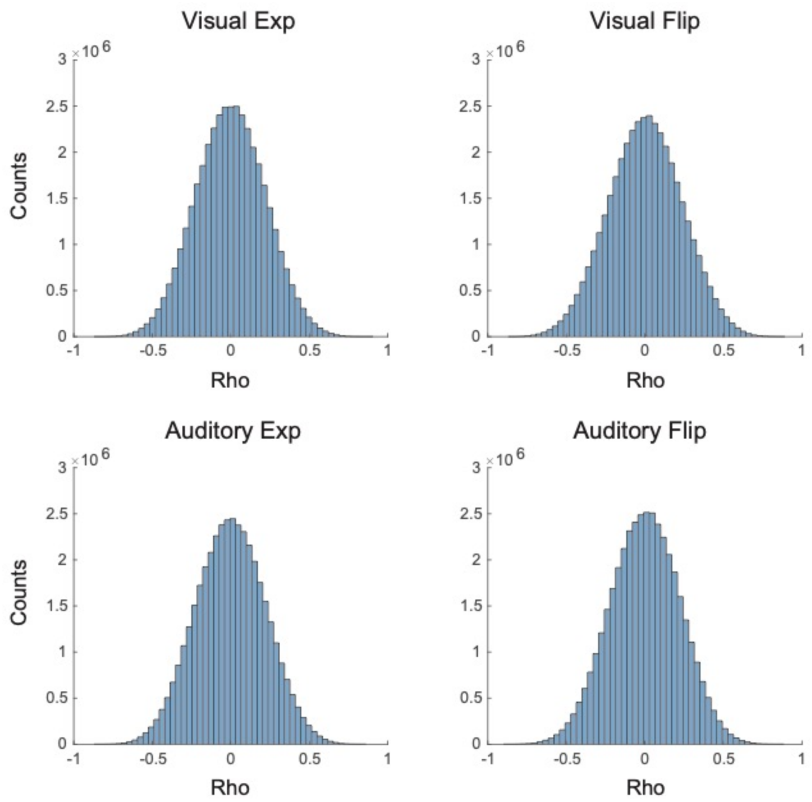


**Fig. S5** Spearman's rho. Rho computed by correlation between source-level ERF of the 'go' cue and RT. ERFs and RT were aggregated within 30 frames (**Methods**).

**Table S3** Time span and location of source clusters computed on Spearman's rho (correlation between ERF of the 'go' cue and RT). Positive clusters are highlighted in red, negative clusters are highlighted in blue.

| | Time span (s) | Location | Time span (s) | Location | Time span (s) | Location |
|---|---|---|---|---|---|---|
| Vis exp | 0.215 - 0.245 | right parietal | 0.220 - 0.250 | left parietal | 0.215 - 0.245 | cerebellum |
| Vis flip | 0.205 - 0.235 | right parietal | 0.215 - 0.245 | left parietal | 0.240 - 0.270 | cerebellum |
| Aud exp | 0.185 - 0.215 | right parietal | 0.210 - 0.240 | left parietal | 0.200 - 0.230 | cerebellum |
| Aud flip | 0.215 - 0.245 | right parietal | 0.195 - 0.225 | left parietal | 0.235 - 0.265 | cerebellum |
| Vis exp | 0.400 - 0.430 | left parietal | 0.455 - 0.485 | right parietal | | |
| Vis flip | 0.395 - 0.425 | left parietal | 0.455 - 0.485 | right parietal | | |
| Aud exp | 0.385 - 0.415 | left parietal | --- | --- | | |
| Aud flip | 0.385 - 0.415 | left parietal | 0.395 - 0.425 | right parietal | | |

**Table S4** P values of source clusters computed on Spearman's rho (correlation between ERF of the 'go' cue and RT). Positive clusters are highlighted in red, negative clusters are highlighted in blue.

| | P value | Location | P value | Location | P value | Location |
|---|---|---|---|---|---|---|
| Vis exp | 0.0005 | right parietal | 0.0085 | left parietal | 0.0075 | cerebellum |
| Vis flip | 0.5 | right parietal | 0.002 | left parietal | 0.0005 | cerebellum |
| Aud exp | 0.17 | right parietal | 0.001 | left parietal | 0.013 | cerebellum |
| Aud flip | 0.042 | right parietal | 0.023 | left parietal | 0.047 | cerebellum |
| Vis exp | 0.018 | left parietal | 0.017 | right parietal | | |
| Vis flip | 0.038 | left parietal | 0.52 | right parietal | | |
| Aud exp | 0.01 | left parietal | --- | --- | | |
| Aud flip | 0.39 | left parietal | 0.72 | right parietal | | |

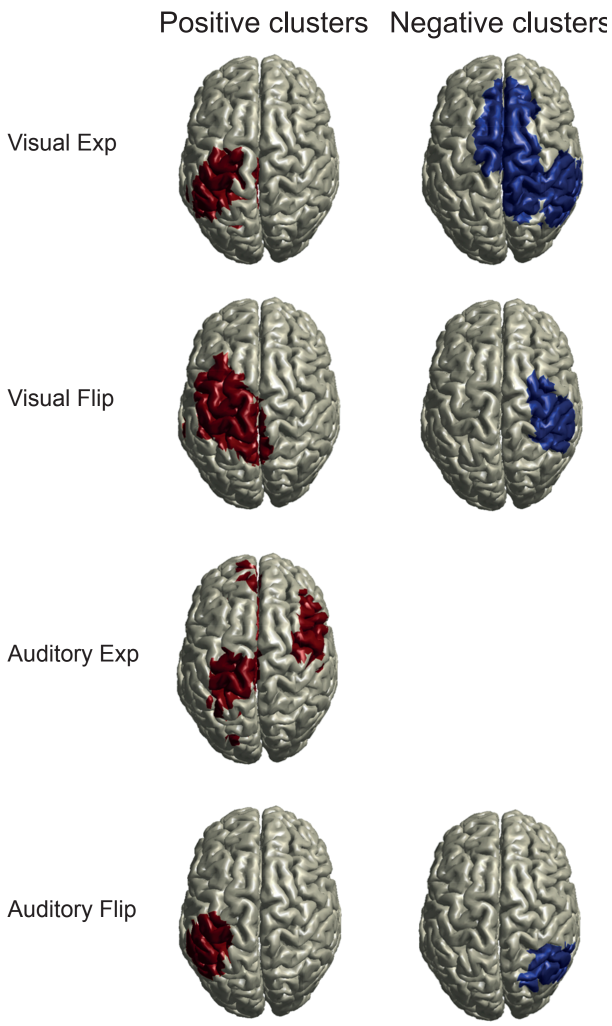


**Fig. S6** Clusters of Spearman's rho around 400 ms post-'go' cue. Rho was computed by correlating source-level representation of 'go' ERF and RT (**Methods**). See **Tbl. 5** for time spans of clusters.

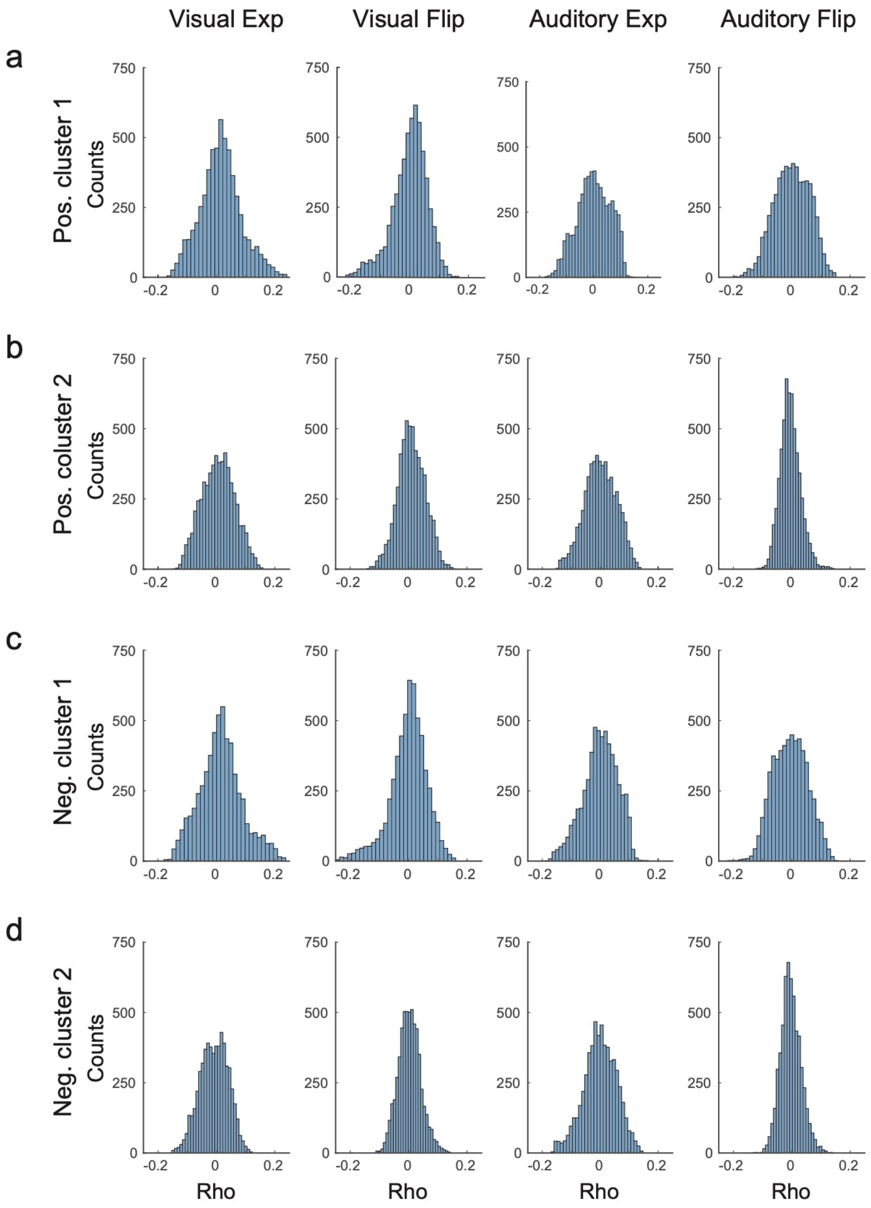


**Fig. S7** Histograms of Spearman's rho. Each plot shows 5798 mean values of rho (averages over the time spans given in **Tbl. 5**). Rho was computed by correlating source-level representation of 'go' ERF and RT (**Methods**).

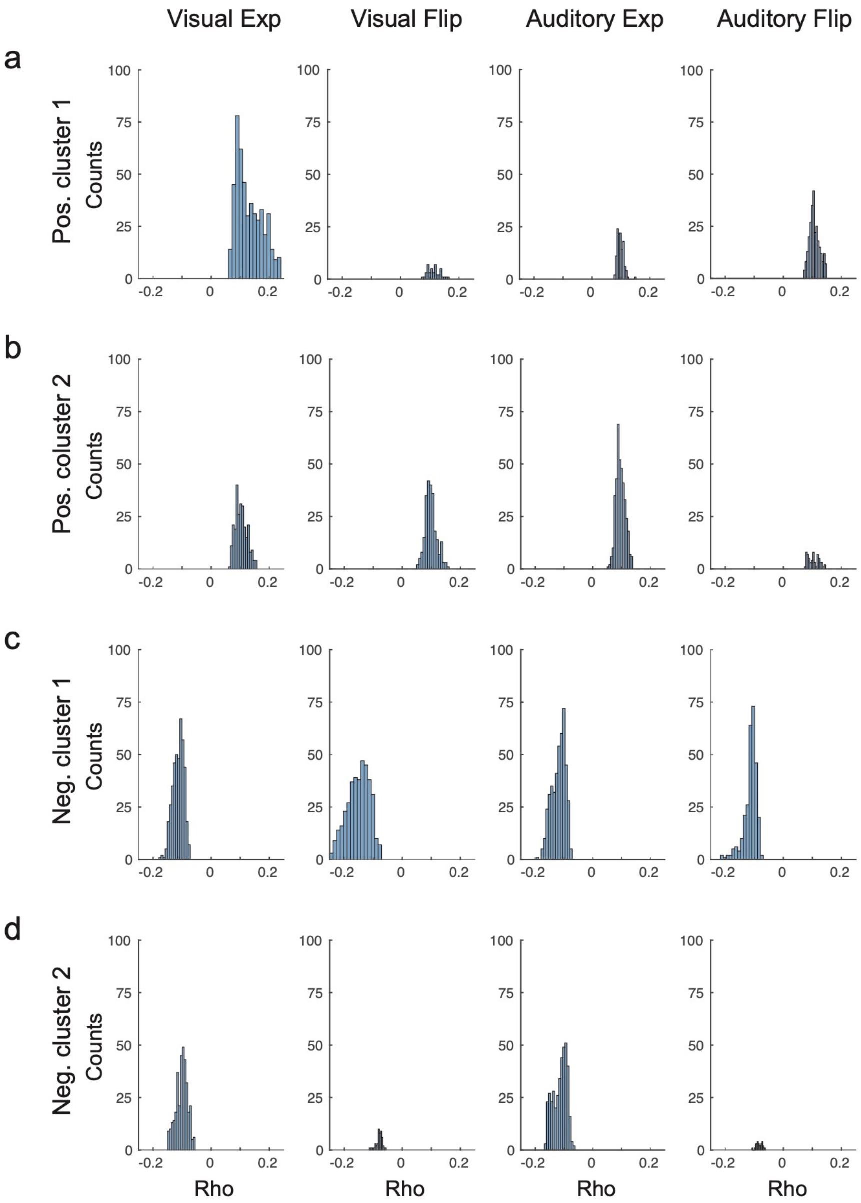


**Fig. S8** Histograms of Spearman's rho per cluster. Rho was computed by correlating source-level representation of 'go' ERF and RT (**Methods**).

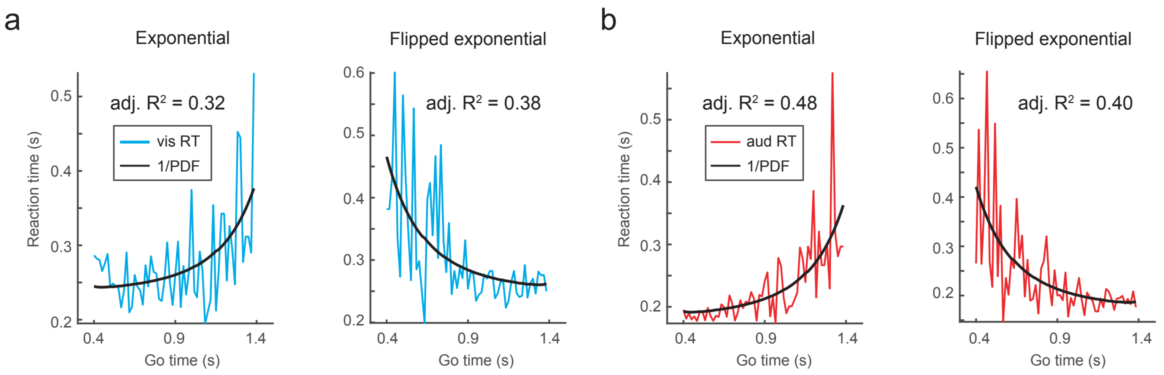


**Fig. S9** 1/PDF fit to a single participant's RT data. **a**) visual conditions, **b**) auditory conditions.

**Table S6** Cluster-based permutation test on Spearman's rho (correlation between sensor-level ERF of the 'go' cue and fit 1/PDF) ERF data baselined using pre-'go' cue baseline (**Methods**).

| Condition | Cue | Regressor | Positive cluster (*P*- value) | | | Negative cluster (*P*- value) | |
|---|---|---|---|---|---|---|---|
| | | | #1 | #2 | #3 | #1 | #2 |
| Vis exp | 'go' | 1/PDF | 0.008 | 0.024 | --- | 0.004 | (0.028) |
| Vis flip | 'go' | 1/PDF | 0.002 | 0.008 | 0.002 | 0.002 | 0.002 |
| Aud exp | 'go' | 1/PDF | 0.004 | 0.008 | --- | 0.002 | 0.01 |
| Aud flip | 'go' | 1/PDF | 0.002 | 0.024 | --- | 0.002 | (0.048) |

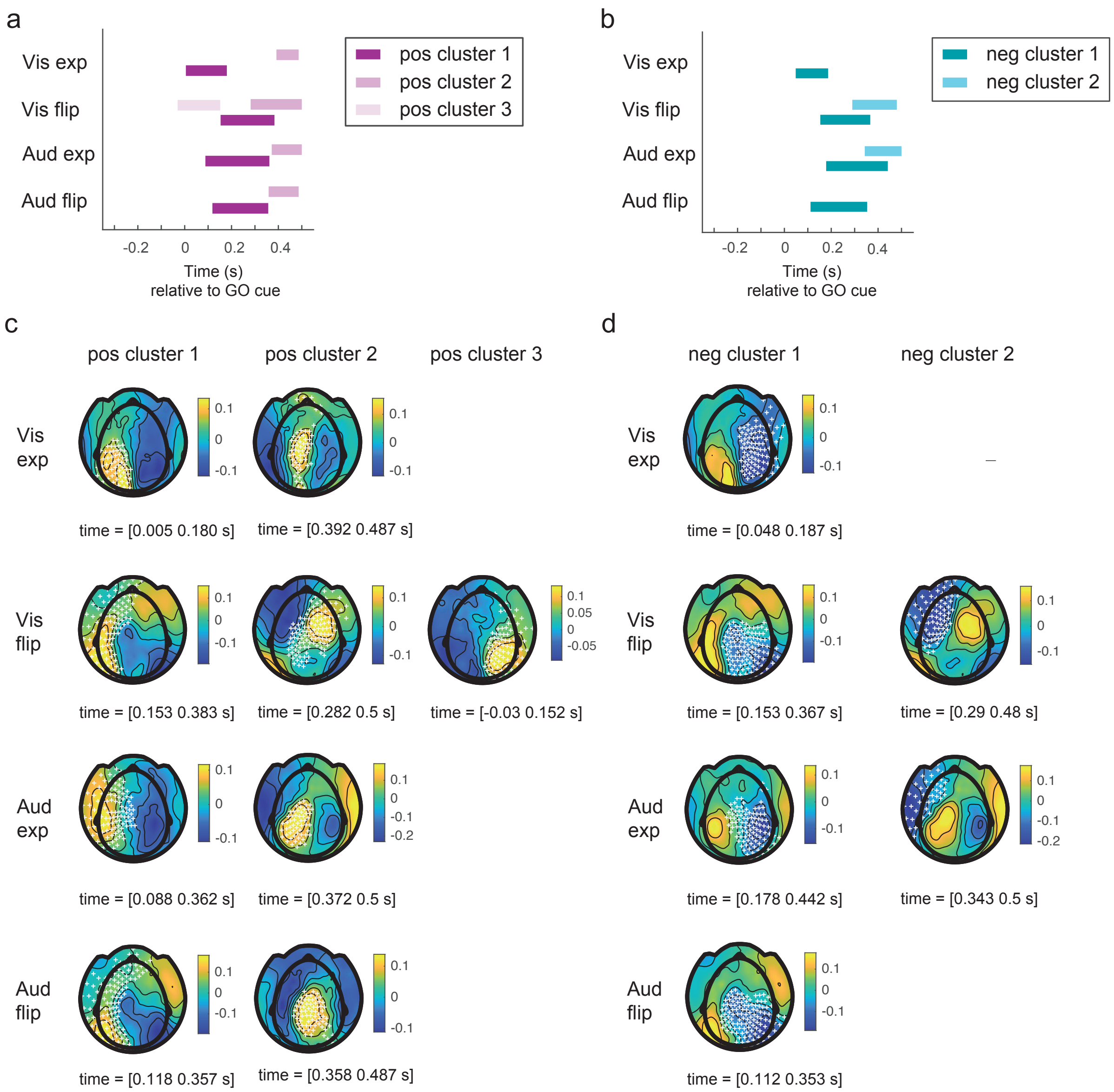


**Fig. S10** Cluster-based permutation test on Spearman's rho (correlation between ERF of the 'go' cue and fit 1/PDF). **a**) Positive clusters **b**) Negative clusters **c**) Topography plots of rho, channels of positive cluster highlighted **d**) Topography plots of rho, channels of negative cluster highlighted**.** A minimum time span of 20 ms was set for a channel to be included in cluster. ERF data baselined using pre-'go' cue baseline (**Methods**).

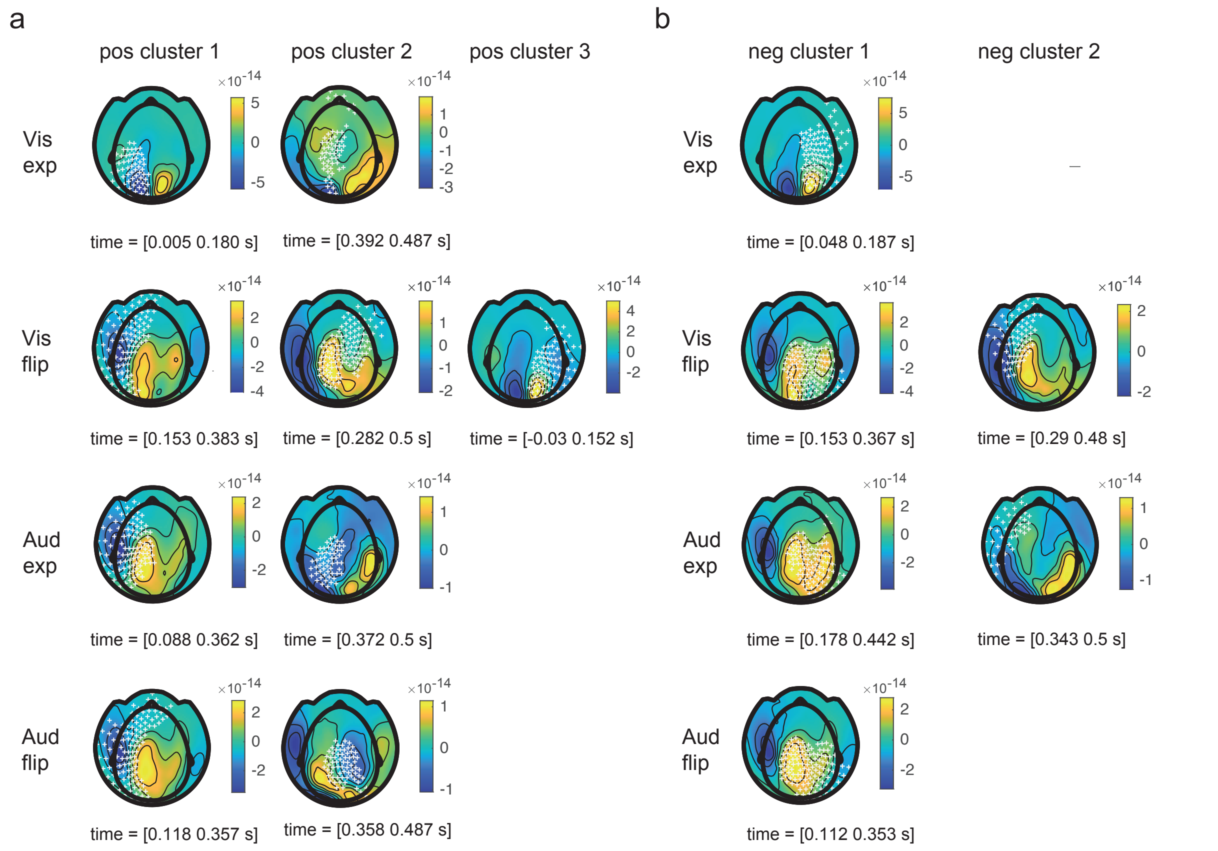


**Fig. S11** Cluster-based permutation test on Spearman's rho (correlation between ERF of the 'go' cue and fit 1/PDF). **a**) Topography plots of 'go' ERF, channels of positive cluster highlighted. **b**) Topography plots of 'go' ERF, channels of negative cluster highlighted. A minimum time span of 20 ms was set for a channel to be included in cluster. ERF data baselined using pre-'go' cue baseline (**Methods**).

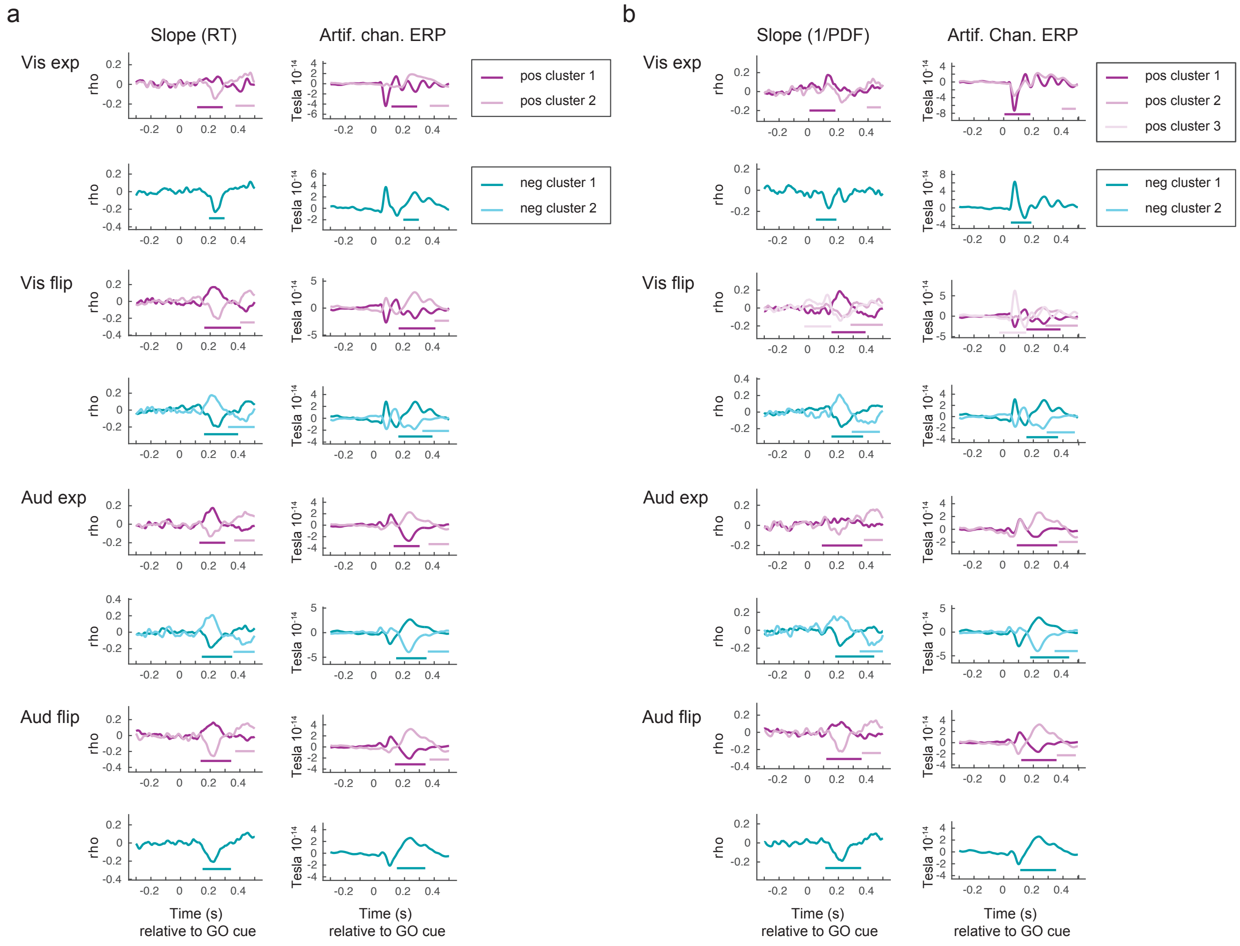


**Fig. S12** Spearman's rho and 'go' cue ERF. **a**) Both rho (correlation between ERF of the 'go' cue and RT) and the 'go' cue ERF were averaged within an artificial channels that comprised all single channels that comprise a cluster and are significant for more than 20 consecutive ms ($P = 0.05$, two-sided). **b**) Same for rho computed by correlation between ERF and fit 1/PDF. Cluster time spans demarkated by horizontal bars. ERF data baselined using pre-'go' cue baseline (**Methods**).

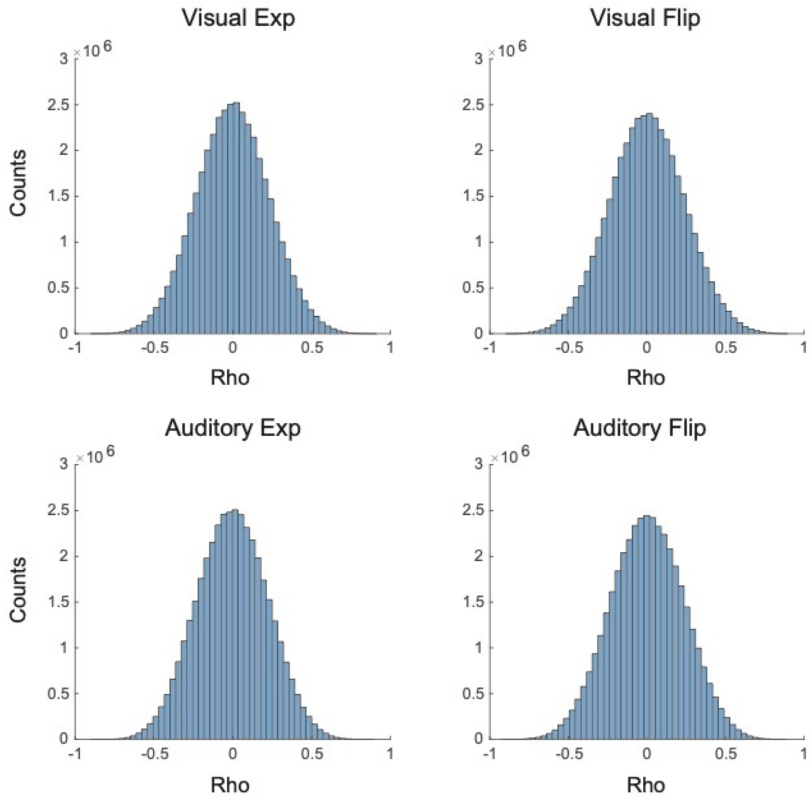


**Fig. 13** Spearman's rho. Rho computed by correlation between source-level ERF of the 'go' cue and fit 1/PDF. ERFs and RT were aggregated within 30 frames (**Methods**).

**Table S7** Time span and location of source clusters computed on Spearman's rho (correlation between ERF of the 'go' cue and fit 1/PDF). Positive clusters are highlighted in red, negative clusters are highlighted in blue.

| | Time span (s) | Location | Time span (s) | Location |
|---|---|---|---|---|
| Vis exp | 0.215 - 0.245 | right parietal | 0.220 - 0.250 | left parietal |
| Vis flip | 0.205 - 0.235 | right parietal | 0.215 - 0.245 | left parietal |
| Aud exp | 0.200 - 0.230 | right parietal | 0.210 - 0.240 | left parietal |
| Aud flip | 0.215 - 0.245 | right parietal | 0.195 - 0.225 | left parietal |

**Table S8** P Values of source clusters (~200 ms) computed on Spearman's rho (correlation between ERF of the 'go' cue and fit 1/PDF). Positive clusters are highlighted in red, negative clusters are highlighted in blue.

| | P value | Location | P Value | Location |
|---|---|---|---|---|
| Vis exp | 0.006 | right parietal | 0.32 | left parietal |
| Vis flip | 0.003 | right parietal | 0.001 | left parietal |
| Aud exp | 0.036 | right parietal | 0.008 | left parietal |
| Aud flip | 0.013 | right parietal | 0.10 | left parietal |

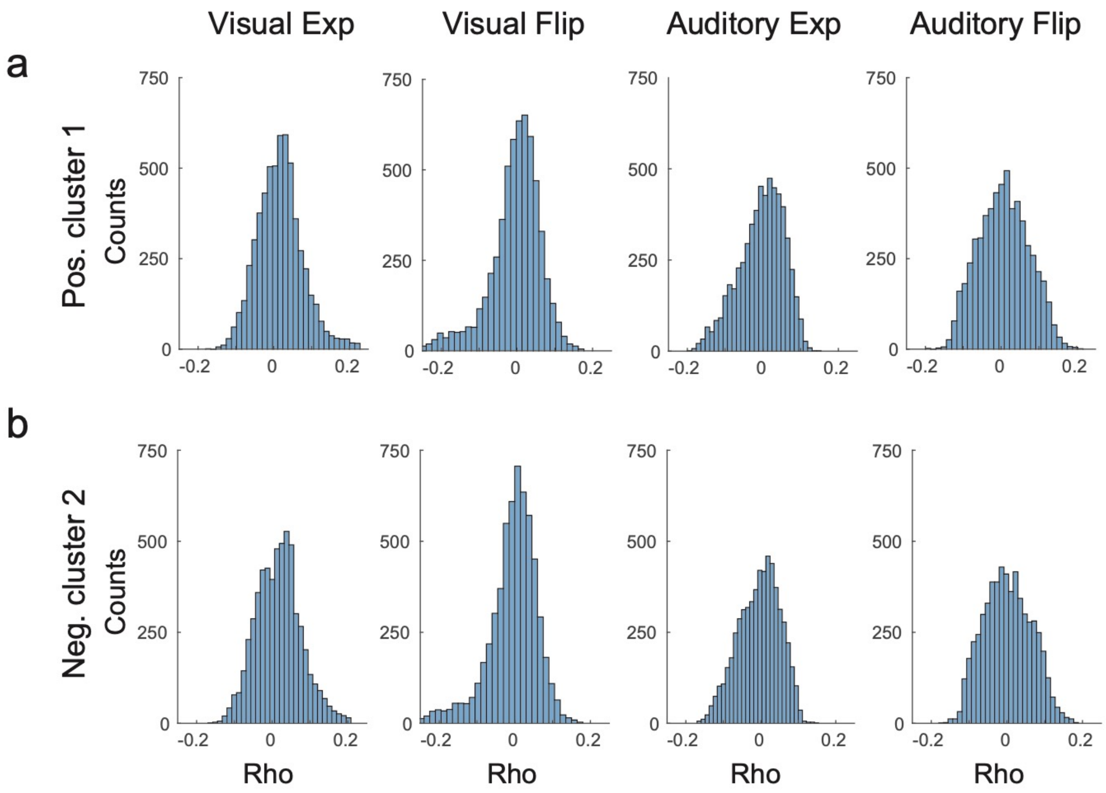


**Fig. S14** Histograms of Spearman's rho. Each plot shows 5798 mean values of rho (averages over the time spans given in **Tbl. 5**). Rho was computed by correlating source-level representation of 'go' ERF and fit 1/PDF (**Methods**).

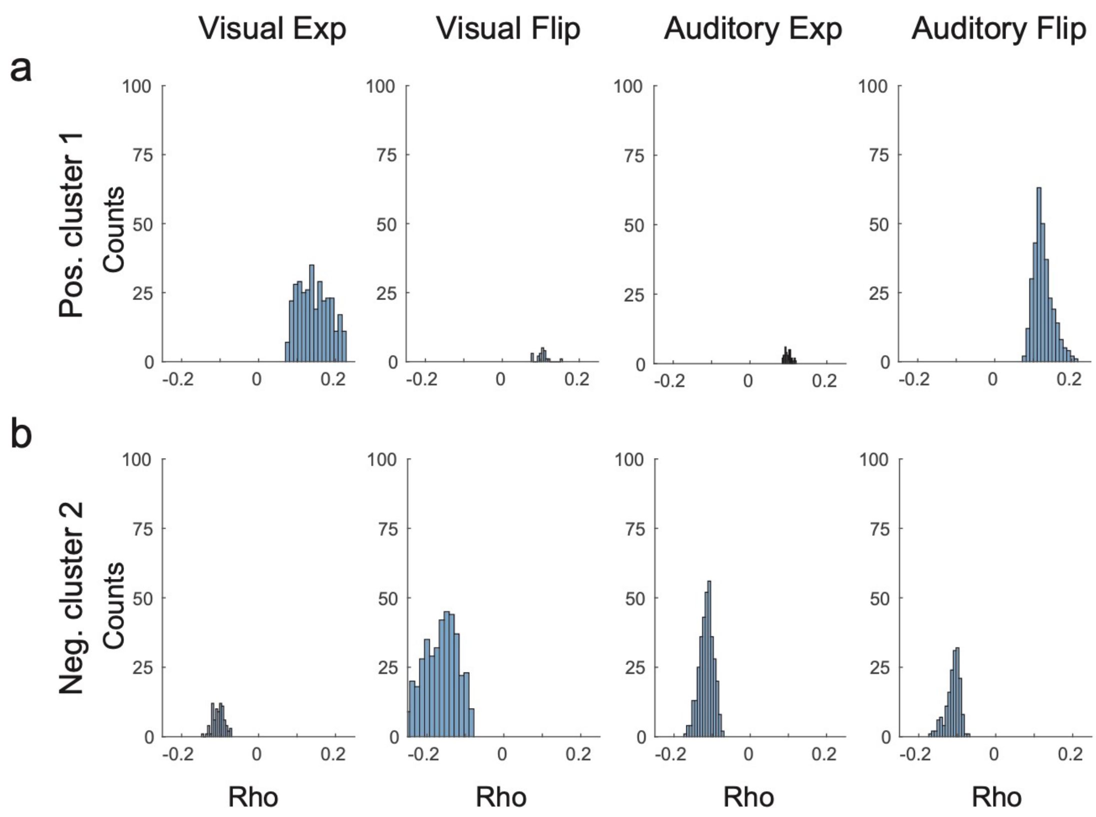


**Fig. S15** Histograms of Spearman's rho per cluster. Rho was computed by correlating source-level representation of 'go' ERF and fit 1/PDF (**Methods**).

### A note on correlation bias

In the ERF analysis, trials were aggregated in 30 frames of equal duration with respect to 'go'-time. The number of trials in each frame varied considerably, as the high probability frames contained many trials, whereas the frames with low event probability contained only few trials. Within each frame, the RT and the MEG were respectively averaged for each subject, condition, channel and time point. So there is an obvious question whether averaging a different number of trials within each frame introduces a bias in the correlation between RT and MEG.

We investigated this possible confound with simulated data resembling the distributions in the actual MEG data. The evoked MEG data had a distribution close to a Gaussian centered around zero. We simulated random data with similar distributions. RT was simulated by the 1/PDF linear model that was presented in the initial behavioral analysis.

The 'go'-time frames had the same distribution of trials as in the actual data. The simulated MEG data was then averaged within frames and the 30 frame values were correlated with the 30 RT simulated values (and the 1/PDF model). As the simulated MEG data was random, also the correlations with simulated RT values was expected to be around zero with no monotonic structure. Any monotonic structure would hint towards an artifact from the averaging of a different number of trials per frame, as the trial number decreases or increases monotonically, depending on the event distribution (exponential-decreasing, flipped-increasing).

This procedure was repeated 10000 times with each time having new random simulated MEG data. The histogram of the resultant 10000 correlation values was plotted for the simulated ERF case (**Fig. 30**) .

In the ERF case, no bias was observed in the distribution of rho, as it was centered round zero. As no bias seems to be present due to the averaging within frames, we proceeded and correlated the within-frames averaged, time-locked MEG data with the within-frames averages of RT.

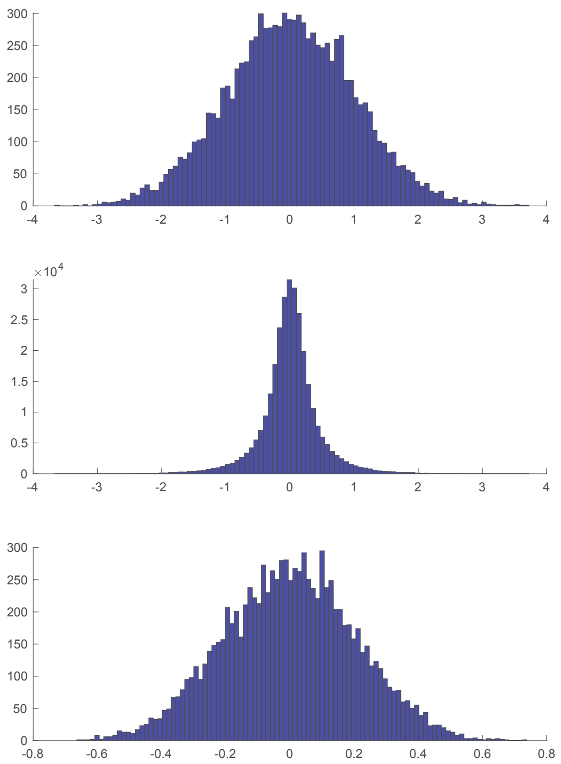


**Fig. S30** Data simulation. Simulated ERF data (top), aggregated within frames (middle), Spearman's rho computed on aggregated simulated ERF data and 1/PDF (bottom).
